# Fast and flexible linear mixed models for genome-wide genetics

**DOI:** 10.1101/373902

**Authors:** Daniel E Runcie, Lorin Crawford

## Abstract

Linear mixed effect models are powerful tools used to account for population structure in genome-wide association studies (GWASs) and estimate the genetic architecture of complex traits. However, fully-specified models are computationally demanding and common simplifications often lead to reduced power or biased inference. We describe Grid-LMM (https://github.com/deruncie/GridLMM), an extendable algorithm for repeatedly fitting complex linear models that account for multiple sources of heterogeneity, such as additive and non-additive genetic variance, spatial heterogeneity, and genotype-environment interactions. Grid-LMM can compute approximate (yet highly accurate) frequentist test statistics or Bayesian posterior summaries at a genome-wide scale in a fraction of the time compared to existing general-purpose methods. We apply Grid-LMM to two types of quantitative genetic analyses. The first is focused on accounting for spatial variability and non-additive genetic variance while scanning for QTL; and the second aims to identify gene expression traits affected by non-additive genetic variation. In both cases, modeling multiple sources of heterogeneity leads to new discoveries.

**Author summary:** The goal of quantitative genetics is to characterize the relationship between genetic variation and variation in quantitative traits such as height, productivity, or disease susceptibility. A statistical method known as the linear mixed effect model has been critical to the development of quantitative genetics. First applied to animal breeding, this model now forms the basis of a wide-range of modern genomic analyses including genome-wide associations, polygenic modeling, and genomic prediction. The same model is also widely used in ecology, evolutionary genetics, social sciences, and many other fields. Mixed models are frequently multi-faceted, which is necessary for accurately modeling data that is generated from complex experimental designs. However, most genomic applications use only the simplest form of linear mixed methods because the computational demands for model fitting can be too great. We develop a flexible approach for fitting linear mixed models to genome scale data that greatly reduces their computational burden and provides flexibility for users to choose the best statistical paradigm for their data analysis. We demonstrate improved accuracy for genetic association tests, increased power to discover causal genetic variants, and the ability to provide accurate summaries of model uncertainty using both simulated and real data examples.

## Introduction

Population stratification, genetic relatedness, ascertainment, and other sources of heterogeneity lead to spurious signals and reduced power in genetic association studies [1–5]. When not properly taken into account, non-additive genetic effects and environmental variation can also bias estimates of heritability, polygenic adaptation, and genetic values in breeding programs [5–8]. Both issues are caused by departures from a key assumption underlying linear models that observations are independent. Non-independent samples lead to a form of pseudo-replication, effectively reducing the true sample size. Linear mixed effect models (LMMs) are widely used to account for non-independent samples in quantitative genetics [9]. The flexibility and interpretability of LMMs make them a dominant statistical tool in much of biological research [9–18].

Random effect terms are used in LMMs to account for specific correlations among observations. Fitting an LMM requires estimating the importance of each random effect, called its variance component. General-purpose tools for this are too slow to be practical for genome-scale datasets with thousands of observations and millions of genetic markers [19]. This lack of scalability is caused primarily by two factors: (i) closed-form solutions of maximum-likelihood (ML or REML) or posterior estimates of the variance components are not available and numerical optimization routines require repeatedly evaluating the likelihood function many times, and (ii) each evaluation of the likelihood requires inverting the covariance matrix of random effects, an operation that scales cubically with the number of observations. Repeating this whole process millions of times quickly becomes infeasible.

To this end, several specialized approaches have been developed to improve the speed of LMMs, including the use of sparse matrix operations [20, 21], spectral decomposition of the random effect covariance matrix [22–26], and Monte Carlo REML [27]. These algorithms are particularly useful when the same random effect structure is used many times. For example, in genome-wide association studies (GWAS), each marker is tested with the same LMM. Similarly, in population-level transcriptomics, eQTLs or variance components are estimated for each of tens-of-thousands of genes expression traits. Fast and exact algorithms for fitting LMMs are limited to the case of only a single (full-rank) random effect term, besides the residuals [22–24]. Recently, approximate learning algorithms have been developed for the scalable extension to multiple random effects [28, 29], but few of these ensure guarantees in terms of estimation accuracy. One strategy applicable to studies with multiple random effects is to estimate the variance components only once, in a model without additional marker effects, and then test each marker either using a score test [30] (which does not produce an effect size estimate), or with a conditional F-test assuming the variance component estimates are fixed [31–34]. Given the “known” variance components, closed-form solutions of all other parameters of an LMM can be found using a rotated version of the simple linear model. Unfortunately, both approximations suffer from reduced power when marker effects are large, intractable posterior inference in a Bayesian analysis, and the inability to be applied to parallel analyses over many traits (like gene expression). Table 1 summarizes these different methods, details their respective computational complexities, and provides relevant references.

**Table 1.**
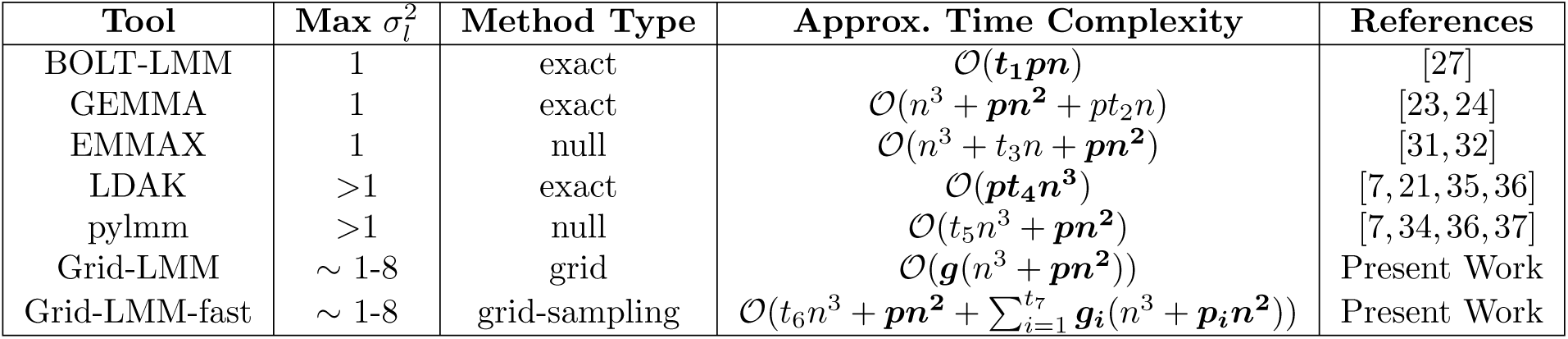
Performance and limitations of reference models for linear mixed model GWAS. The time complexity of each algorithm is approximate, assuming a model with only a single marker effect and no other fixed effects. Here, *l* is used to index full-rank random effects; 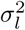 are variance component parameters; *n* is the number of observations; and *p* is number of markers to test. Denote *t*_1_ *… t*_7_ to represent the number of iterations needed for convergence, which is expected to vary among methods (particularly for the iterations of grid search in `Grid-LMM-fast`), and may vary across markers. The terms *g* and *g_i_* are grid sizes for the `Grid-LMM` methods (i.e. the number of grid vertices that must be evaluated). Lastly, *p_i_* is the number of markers that need to be tested in iteration of *i* ϵ {1…*t*_7_} of the `Grid-LMM-fast` method. The rate limiting terms in common GWAS applications (where *p* ≫ *n*) are in bold. “Method Type” describes the estimation of 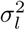. “Exact” means effectively exact, up to machine precision and subject to possible convergence of algorithms to local maxima. “Null” means estimation of parameters under the null model with no marker effects. References list additional methods that are approximately equivalent to the given model classes.

`Grid-LMM` takes a different approach to fitting LMMs: rather than directly optimizing the variance components separately for each test, we define a grid spanning all valid values of the variance components and fit simple linear models at each grid location. Each evaluation involves a single Cholesky decomposition of the random effect covariance matrix, which is then reused to calculate closed-form ML solutions or Bayesian posterior summaries (under a specific conjugate prior; see S1 Supplementary Methods) for all separate tests. This results in dramatic time-savings in typical GWAS settings (see S1 Fig). After repeating these calculations across the whole grid, we select the highest ML (or REML) score for each marker to compute approximate likelihood ratio and Wald tests [32, 38], or analogously derive posterior distributions and Bayes factors by summing appropriate statistics across the grid. The `Grid-LMM` approach relies on a re-parameterization of the typical LMM framework from individual variance components 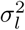 to variance component proportions 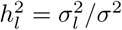, where *σ*^2^ without the subscript denotes the total sum of all variance components (including the residual). Since the variance components must be non-negative, their proportions are restricted to the unit interval [0, 1] and sum to 1, forming a simplex. Therefore, a finite grid can span all valid parameter values. While the size of the grid increases rapidly with the number of random effects, for a small number of random effects (~1-8) and a moderate grid resolution, the size of the grid remains tiny relative to the number of models in a typical GWAS. As we show below, highly accurate test statistics are achieved even with a coarse grid in most reasonable situations, and further improvements to efficiency are possible by using heuristics to adaptively sample the grid or reduce the number of grid locations computed for the majority of tests. This strategy of conditioning on variance components over a grid and then combining solutions can be applied to many other tools in quantitative genetics including set tests for rare variants [39, 40], whole-genome regression models such as LASSO and elastic net [41, 42], and QTL mapping in controlled crosses [43].

In the following sections, we demonstrate the accuracy and advantages of the `Grid-LMM` approach using a simulation study and two real genome-wide quantitative genetics examples. The first is for GWAS, where tens-to-hundreds of thousands of markers are individually tested for associations with a single phenotype. The second is for gene expression, where thousands of traits are each tested for non-additive genetic variance. In both cases, the same random effect structure is used for each model. While approximate, the test-statistics and posterior quantities calculated by `Grid-LMM` are accurate and improve power relative to other approximation methods — all while maintaining dramatically reduced computational burdens than the direct approach. Full derivations of the model and useful heuristics are described in detail in the Methods.

## Results

As a first case-study, we used `Grid-LMM` to perform two types of genome-wide association studies (GWAS) that benefit from modeling multiple random effects: (1) the study of gene-environment interactions, and (2) detecting associations for phenotypes driven by non-additive genetic variation or spatial variation. In both cases, there are multiple sources of covariation among observations that can inflate test statistics and bias estimates of heritability if not appropriately accounted for by using mixed models.

As an example of a GWAS with gene-environment interactions, we analyzed data on flowering times for *Arabidopsis thaliana* [44]. First, we benchmarked results from standard LMM methodologies to confirm that `Grid-LMM` association tests are accurate. We ran `Grid-LMM` on both a fine-grained grid with a small step size of 0.01 *h*^2^-units, and a larger step size of 0.1 *h*^2^-units, to test for associations between 216,130 single nucleotide polymorphisms (SNPs) and flowering times of 194 accessions measured at 10C (i.e. in a constant environment). We compared the Wald-test *p*-values computed by both `Grid-LMM` models to *p*-values computed using the exact method `GEMMA` [24], and the approximate method `EMMAX` [32]. Each method was applied as an LMM with only the additive relationship matrix as its single random effect. Grid-LMM *p*-values computed using the finer grid size (i.e. 0.01 *h*^2^-units) were almost identical to those of `GEMMA`, and even *p*-values computed with the larger step size (i.e. 0.1 *h*^2^-units) were more accurate than those resulting from `EMMAX`. There were particularly noticeable differences in performance for markers with larger scores, which were strongly underestimated by `EMMAX` since its approximations of *h*^2^ are made strictly under the null model (Fig 1a). This pattern held true across all of the available 107 *Arabidopsis thaliana* phenotypes (see S2 Fig). However, despite these results, we do not advocate `Grid-LMM` in this setting; GEMMA (and similar methods) provides an exact test and is more computationally efficient. The real advantage of `Grid-LMM` is its ability to effectively model two (or more) random effects.

**Fig 1.**
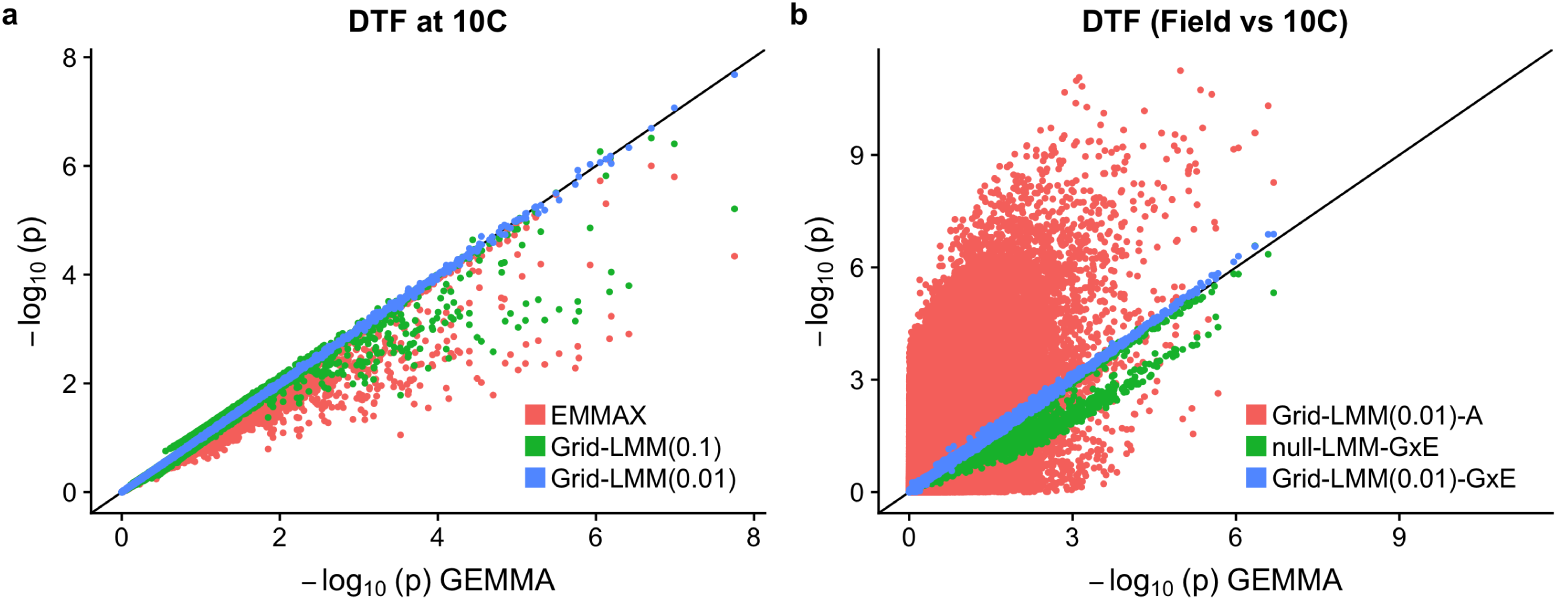
Comparisons of GWAS results for *Arabidopsis* flowering time. (**a**) Results for days-to-flower (DTF) at 10C. Compared are `EMMAX` and `Grid-LMM` with grid sizes 0.1 and 0.01, respectively. The exact method `GEMMA` is treated as a baseline. Each method was applied to the same LMM with only the additive relationship matrix as a random effect, and *p*-values were computed using the Wald test. (**b**) GWAS for the gene-environment (G × E) interactions effects on DTF between the constant 10C and field conditions. `GEMMA` results are given for the plasticity of each accession (i.e. the difference DTF_Field_ - DTF_10C_), and fit with a single random effect of the additive relationship matrix. The other three methods consider the full data with two observations per accession. Namely, `Grid-LMM-A` fit a “standard” GWAS with a grid step size 0.01, but with two random effects — the additive relationship matrix and an *iid* line effect. The other two models, `null-LMM-G × E` and `Grid-LMM-G × E`, fit three random effects — the additive relationship matrix, an *iid* line effect, and a G × E-additive relationship matrix representing the background covariation in gene-environment interactions among accessions.

To demonstrate this advantage, we tested for gene-environment (G × E) interaction effects on flowering time. Here, we combined flowering time data from two conditions: constant 10C (as described above) and in the field. We limited our analysis to the 175 accessions that were grown under both conditions, yielding a total of *n* = 350 observations. When observations come from different environments, we might expect phenotypes to cluster for at least two reasons: (i) the sharing of alleles with constant effects across environments due to relatedness and population structure, commonly modeled with the additive genomic relationship matrix (**A**) as a random effect, and (ii) the sharing of alleles with effects that differ among environments (or are specific to one environment). Previous work has shown that, when testing for G × E effects with GWAS, modeling the second source of covariance by using a second random effect can prevent spurious signals and increase power [34]. The same result is replicated in this setting using simulations (see S3 Fig and S4 Fig).

Here, we calculated G × E *p*-values for each genetic marker using `Grid-LMM` (grid step size = 0.01 *h*^2^ units) on the full dataset using both the **A** and the G × E relationship matrices as two random effects. These are compared to *p*-values from (i) an LMM which ignores the G × E covariance and only considers genetic similarity, and (ii) a model similar to `pylmm` [34] which does consider both random effects but estimates variances components from the null model (this is referred to as `null-LMM-G × E` below). For each model, we included an additional random effect to account for the repetition of accessions which could induce additional covariance among observations. In this dataset, a simpler approach to testing for G × E was also available: test for associations between markers and the *plasticity* (i.e. the difference in flowering time between the field and 10C) of each accession, which requires only a single random effect and can be fit with `GEMMA`. This simple approach is only possible because every genotype was measured in both environments. Nonetheless, it is expected to produce identical tests to the full three-random-effect model and, therefore, serves as a viable benchmark for the `Grid-LMM` results. We used the plasticity GWAS approach as a baseline to compare the three models that use the raw data directly (Fig 1b). As expected, ignoring the G × E covariance leads to greatly inflated tests for the majority of markers. `Grid-LMM` replicated `GEMMA`’s plasticity *p*-values almost exactly when run with three random effects; alternatively, a portion of the `null-LMM-G × E`’s tests were deflated, implying lower power. The full analysis using `Grid-LMM` with three random effects took 26 minutes. Fitting the same model for all 216,130 markers directly using an exact method (e.g. `LDAK` [7]) would take approximately 6 hours (about 14 × longer). Note that `LDAK` is not designed for re-estimating variance components for each SNP in multiple-random-effect models and so, to conduct time comparisons, we simply ran the program multiple times. This requires `LDAK` to re-load the covariance matrices for each marker. However, by controlling the maximum number of iterations in the `LDAK` optimization engine, we estimate that a maximum ≈ 33% of the running time for these models is due to data import, with the remainder being due to the numerical calculations.

Even when all individuals are measured in the same environment, the typical additive relationship matrix may not account for all sources of covariation. In particular, spatial variation and the sharing of non-additive alleles may also induce covariance, lead to reduced power, and/or result in inflated tests in GWAS [45]. As an illustrative example, we tested for genetic association between 10,075 bi-allelic autosomal markers and body mass among 1,814 heterogenous stock mice from the Wellcome Trust Centre for Human Genetics (WTCHG) [46]. We first computed additive and pairwise-epistatic relationship matrices, as well as a spatial-environmental covariance matrix based on the 523 different cages used in the experiment. Next, we compared *p*-values derived from an LMM considering only the additive relationship matrix (as in a typical GWAS) to those calculated by `Grid-LMM` using all three relationship matrices (Fig 2a-d). Using the three-random effect model, we identified associations on two chromosomes (numbers 4 and 11) that were not apparent when just using the additive relationship matrix as a random effect. Both loci have been previously identified as size-associated QTL (see Table S1 Table) [47, 48]. Genomic control values for the two models were both close to one (A-only = 0.975, A+E+Cage = 0.974) (Fig 2b,d). The three-random effect model took 8.5 minutes to fit using `Grid-LMM`, while a full analysis on all 10,346 markers would have taken approximately 10 hours with `LDAK` (more than 100 × longer, of which we estimate a maximum of ≈10% is spent on reading in data). Extrapolating to a consortium sized genome-wide analysis with 1 million markers would take ≈ 14 hours using `Grid-LMM`, as opposed to 40 days using `LDAK`. We see larger performance gains in the mouse dataset compared to the *Arabidopsis* dataset because of the larger sample size (*n* =1,814 vs. 350). `LDAK` (and other exact general-purpose REML methods) is dominated by matrix inversions which scale with *n*^3^, while `Grid-LMM` is dominated by matrix-vector multiplications which scale with *n*^2^ (again see Table 1). The performance advantage of `Grid-LMM` will increase even more for datasets with more individuals.

**Fig 2.**
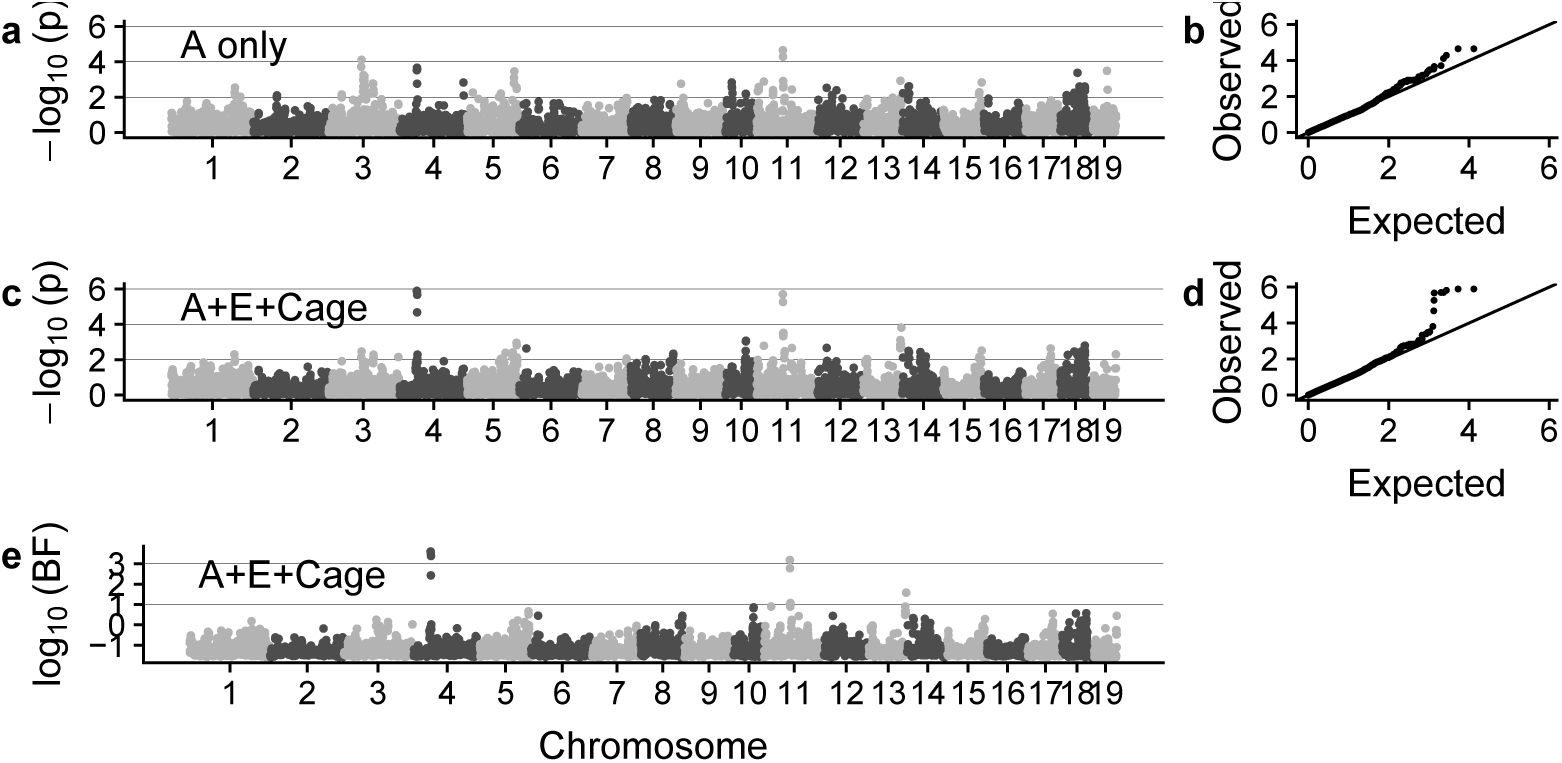
GWAS results for the body weight trait of the Wellcome Trust heterogenous stock mice. `Grid-LMM` was run using a grid size of 0.01 *h*^2^-units for each model. (**a**-**b**) Wald test *p*-values from a model with only the additive relationship matrix (**A**) as a random effect, organized by chromosome and compared to a uniform distribution via a QQ-plot. (**c**-**d**) Wald test *p*-values from a model with three random effects: the additive, epistatic (**E**), and spatial (**Cage**) relationship matrices. (**e**) Approximate Bayes factors calculated from the three random effect model, assuming a standard normal N(0, 1) prior for the marker effects and a uniform prior on the grid of variance component proportions.

To demonstrate the flexibility of `Grid-LMM` for GWAS, we also ran an analysis on the mice body weight trait using Bayesian inference and calculated Bayes Factors by comparing the models for each marker to the null model assuming no genetic effects. Here, we used a uniform prior over the grid of variance component proportions and assumed a standard normal prior for the marker effect sizes. In this setting, the Bayes Factors were highly correlated with the frequentist *p*-values — they also highlight the same associations on chromosomes 4 and 11 (Fig 2e). In general, Bayes Factors provide additional flexibility for incorporating prior information on individual markers or even combining results among multiple studies [49, 50]. The full Bayesian analysis for `Grid-LMM` took 15.5 minutes, just 7 minutes longer than the frequentist analysis, making it practical for genome-wide studies.

As a third case-study, we used `Grid-LMM` to estimate the additive and pairwise-epistatic variance components for 20,178 gene expression traits measured on 681 *Arabidopsis* accessions from the 1001 Genomes Project [51]. Using a grid-size of 0.05 *h*^2^ units, we estimated the magnitude of each variance component by REML (Fig 3a). The whole analysis took ≈ 6 minutes. Finding REML solutions for the same two-random effect model, on each of the traits separately, using the exact method `LDAK` took ≈ 90 minutes (of which ≈ 20% was due to data import). Grid-LMM variance component estimates replicated those of `LDAK` accurately, but with less precision due to the coarse grid size (see S5 Fig). Notably, for many genes, the proportion of variance in expression attributed to additive variance dropped considerably when the epistatic variance was also modeled (see S6 Fig). Therefore, including multiple random effects can have significant impact on downstream conclusions for many traits.

**Fig 3.**
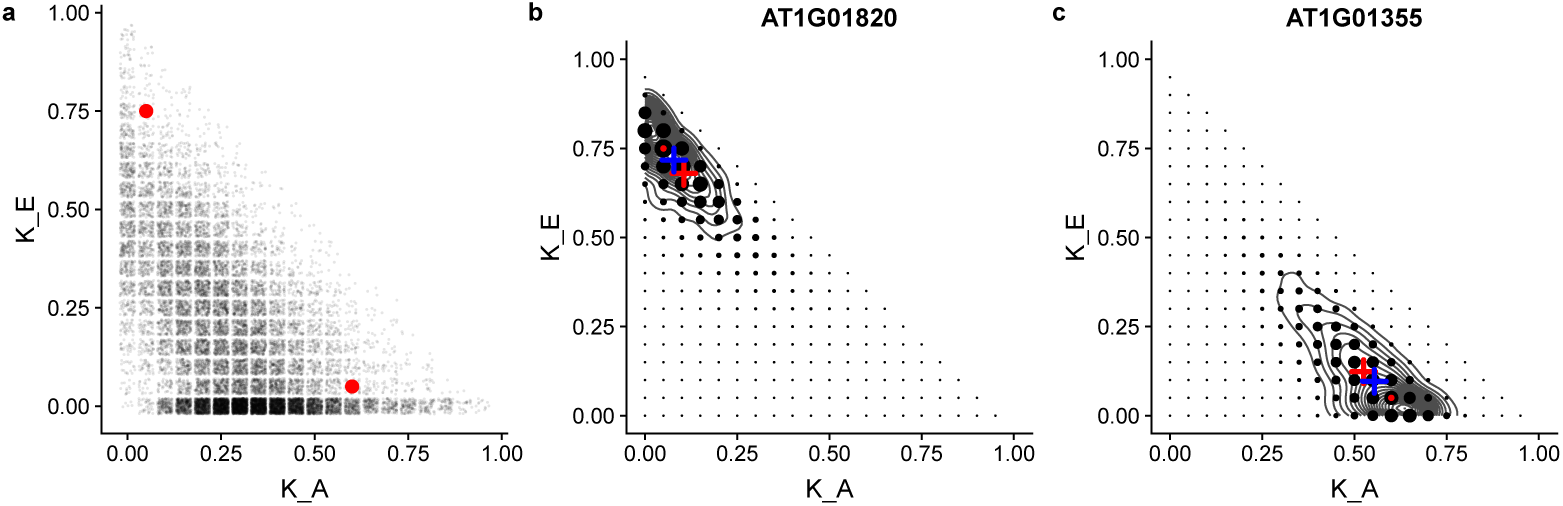
Variance component estimates for *Arabidopsis* gene exprssion traits. `Grid-LMM` was fit using a grid size of 0.05 *h*^2^-units for each of 20,178 genes. (**a**) REML estimates of the variance components proportions for the additive (**K**_*A*_) and pairwise-epistatic (**K**_*E*_) genetic variances. Estimates are jittered for clarity. (**b**-**c**) Posterior distributions of variance component proportions for two example genes, highlighted with red dots in (**a**). The area of each point is proportional to the posterior mass at that combination of the two variance components. A uniform prior over the grid was assumed, and the intercept was assigned a Gaussian prior with infinite variance. The grey contours show the posterior density as estimated by `rstan` using half-Student-t(3,0,10) priors on each standard deviation parameter. In each plot, the red dot shows the REML estimates. The red cross is the posterior mean as estimated by `Grid-LMM`. The blue cross shows the posterior mean as estimated by `rstan`.

Even with this relatively large sample size for a population-level gene expression dataset, considerable uncertainty remains in the estimated variance components. The point-estimates calculated by REML do not capture this uncertainty and estimating confidence intervals for the variance components using REML is difficult. The full `Grid-LMM` approach can be used to calculate posterior distributions for each variance component with little additional cost — note that MCMC sampling is not needed because a reasonably-sized grid can span all valid values of each variance component proportion (see Methods). Using this approach, we identify 8,585 genes with a posterior probability of non-zero epistatic variance greater than 90%, and 28 more genes with a posterior mean epistatic variance component greater than 80%. For two example genes, we show that the fitted posterior distributions are similar to those estimated via MCMC using rstan [52, 53] (see Fig 3b-c). The `rstan` analyses took ≈ 20 hours per gene to generate an effective sample size of ≈ 200 − 400 for the variance component parameters. Therefore, posterior inference by MCMC for all 20,178 genes would take about 50 years of computational time.

Now that we have demonstrated in real data examples that `Grid-LMM` is accurate, fast, and expands the range of genetic analyses that are computationally feasible, we turn to the question of scalability. Specifically, we assess whether `Grid-LMM` is sufficiently accurate for the much larger sample sizes commonly used in modern human GWAS’s. There are no conceptual hurdles to implementing `Grid-LMM` for studies with tens-to-hundreds of thousands of samples and the improvement in time, relative to a direct mixed modeling approach, should increase dramatically (see S1 Figa). Unfortunately, total computational time and memory requirements will grow significantly as well (see S1 Figb). For example, storing a single Cholesky decomposition of the random effect covariance matrix for 100,000 samples would require approximately 80 Gb RAM and would take approximately two days to compute. This contrasts with `BOLT-LMM` which can run a GWAS analysis of this size in less than an hour, while using less than 10 Gb RAM [27]. However, `BOLT-LMM` is restricted to a single random effect and uses a “null-LMM” approach for estimating variance component parameters as part of a two-step analysis.

To test if `Grid-LMM`’s accuracy changes with larger sample sizes, we artificially increased and decreased the sample size of the WTCHG mouse dataset by a factor of 5, simulated phenotypic data based on a randomly selected marker and the same three random effects used in the WTCHG analysis above. We then compared the marker’s p-values calculated with the exact mixed model (`Exact-LMM`) to those calculated using `Grid-LMM` with two resolutions: 0.1 and 0.01 *h*^2^ units. As a baseline, we also calculated p-values with the two-step method that estimates variance components under the null (`null-LMM`). See Methods for details on the simulations. For the `Grid-LMM` tests, we assumed that the nearest grid vertices were exactly centered around the variance component estimate from the exact mixed model. This represented a “worse-case scenario” for `Grid-LMM`.

As a function of the variance contributed by the tested marker, the mean relative difference in p-values among the four methods was approximately constant across the three sample sizes (Fig 4a). There were large differences when the marker effect was large, diminishing to no difference for small effects. `Grid-LMM`(0.01) was barely distinguishable from the `Exact-LMM` across all sample sizes and marker effect sizes. Mean (−log_10_) p-values from `Grid-LMM`(0.1) and `null-LMM` were similar to Exact-LMM for small effect sizes, but `Grid-LMM`(0.1) was closer to `Exact-LMM` for large effect sizes. This is expected because most randomly selected markers are correlated with the dominant eigenvectors of the additive relationship matrix; hence, large effect markers will affect the variance attributed to the random effect, but small effect markers will not.

**Fig 4.**
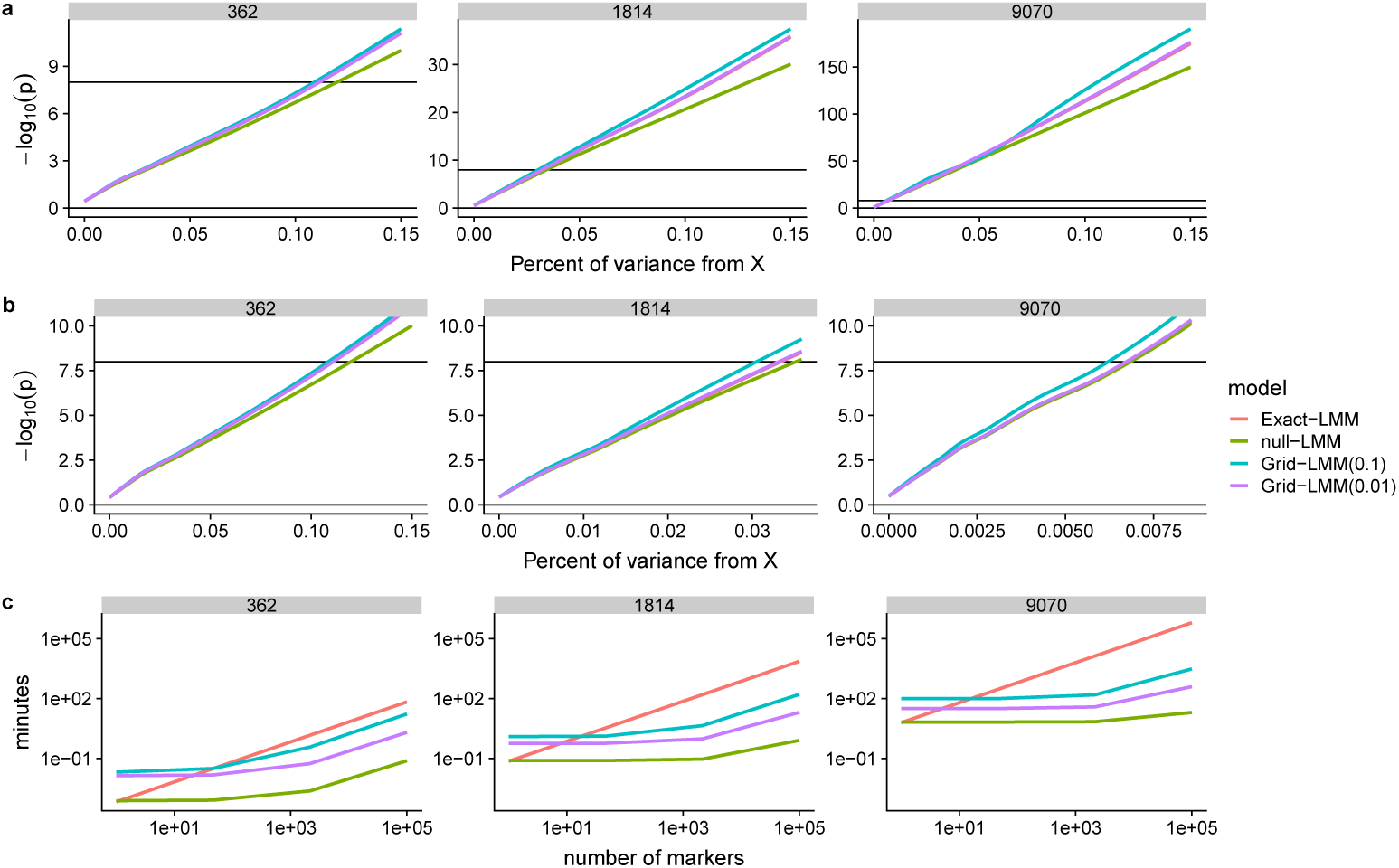
Grid-LMM is accurate but less important at larger sample sizes. We used a simulation study to assess the accuracy of the `GridLMM` approximations for GWAS at larger sample sizes. Data were simulated for three sample sizes (*n* = 362, 1,814 and 9,070), based on the WTCHG mice genotype data. Each simulation was parameterized such that a single randomly chosen marker explained a defined percentage of the total variance (i.e. 0 - 0.15%), while the remainder of variance was simulated based on the additive, epistatic, and cage variance components estimated from the body weight trait. Each simulation was repeated 300 times with different causal markers. The −log_10_ (p)-values were calculated with 4 methods: (i) `Exact-LMM`, an exact LMM implementation; (ii) `null-LMM`, a two-step method with the variance component percentage estimated under the null model; and (iii)-(iv) `Grid-LMM` with grid sizes of 0.1 and 0.01 *h*^2^ units, respectively. For the `Grid-LMM` tests, we assumed that the grid was exactly centered around the variance component estimate from the exact mixed model — a “worse-case scenario” for `Grid-LMM`. (**a**) curves were fit using the `geom_smooth` function from the `ggplot2` package [54]. The horizontal line is at −log_10_(p) = 8 and represents a typical genome-wide significance level. (**b**) The same curves as in the previous row, but zoomed in to the approximate range of marker effects with power ≪ 1 for each sample size. Note that the y-axes are different for each sample size in (**a**), and the x-axes are different for each sample size in (**b**). The `Exact-LMM` curve is hidden by the `Grid-LMM`(0.01) curve in each panel. (**c**) Estimated running times for a GWAS using each method, plotted as a function of the number of markers. The `Exact-LMM` time is computed based on running `LDAK` separately for each marker. The time for `null-LMM` includes a single run of `LDAK` under the null model, and also incorporates the separate tests for each marker conditioning on “known” variance components. The time for `Grid-LMM`(0.1) includes tests for each marker at each of 220 grid vertices constituting a full grid over three variance components. Lastly, `Grid-LMM`(0.01) uses the “fast” heuristic, assuming only one marker has a non-zero effect with an effect size approximately at the power threshold given the sample size (≈ 0.1 for *n* = 362, ≈ 0.035 for *n* = 1814, ≈ 0.006 for *n* = 9070). Lines depict the mean run times based on 10 replications for each calculation.

While the relative change in (−log_10_) p-values is approximately constant, the range of effect sizes where the approximate methods have a negative impact on power changes across sample sizes. Assuming a genome-wide significance threshold of −log_10_(p) = 8 in this dataset, even the `null-LMM` method will consistently declare any marker with an effect size > ≈ 0.02% of the total variance as significant for sample sizes ≈ 10, 000. If we focus specifically on the range of effect sizes where the difference among methods may have an impact on power, the relative performance of the approximate methods do change. At the smallest sample size (i.e. *n* = 362), mean −log_10_(p)-values of `Grid-LMM`(0.1) were closer to those of `Exact-LMM` than those of `null-LMM`. At the medium and large sample sizes (*n* = 1814 and *n* = 9070), the mean −log_10_(p)-values from `null-LMM` were more accurate than those from `Grid-LMM`(0.1). However, the results of the finer `Grid-LMM`(0.01) model remained nearly indistinguishable from those of `Exact-LMM`, irregardless of the number of samples. Note that when using the `Grid-LMM`-fast heuristic, `Grid-LMM` will never perform worse than `null-LMM` because we peg the grid to the variance component estimate under the null model.

Fig 4c compares the estimated running times of GWAS analyses with different numbers of markers for each of the three sample sizes. With *>* 100 markers, results for the `Grid-LMM` methods were intermediate between `Exact-LMM` and `null-LMM`, with the advantage over `Exact-LMM` increasing for large sample sizes. At all sample sizes, `Grid-LMM`(0.1) is linearly slower than `null-LMM` since it effectively requires running `null-LMM` at each grid vertex. At small and intermediate sample sizes this speed penalty is justified by increased power. At large sample sizes `null-LMM` is just as accurate for effect sizes relevant to power, so `Grid-LMM` is not needed (Fig 4b).

## Discussion

Grid-LMM addresses a central obstacle to the practical use of linear mixed models: the computational time needed to find optimal solutions for variance components. Our key observation is that for many common quantitative genetics analyses, optimizing variance component parameters to high precision is not necessary. When sample sizes are large, statistical power will be sufficient to detect associations even under the approximate `null-LMM` methods such as `EMMAX` [32], `pylmm` [34], or BOLT-LMM [27]. However, when sample sizes are more limited, as in the examples we have shown here, the closer approximations achieved by `Grid-LMM` can increase power without greatly increasing computational requirements. Such sample sizes are common in model systems genetics, evolutionary biology, agricultural biology, as well as in eQTL studies. From a Bayesian perspective, the posterior distribution of variance components tends to be broad even with large sample sizes, and coarse approximations can be sufficient given the uncertainty in their true values [55]. In GWAS applications, magnitudes of variance components are of less interest than the accuracy of test statistics for the (fixed) SNP effects, and we show that these are sufficiently accurate even with approximate variance component proportions.

The advantage to relaxing the need for perfect variance component solutions is a vast reduction in both computational time and algorithmic complexity. This reduces the time required for a typical GWAS sized dataset with two-or-more random effects from days to hours, and provides a framework for applying LMMs to even more powerful statistical tools [56–58]. We optimize variance component estimation with a grid search, the simplest type of optimization algorithms. At each grid vertex, after conditioning on (relative) variance component values, the LMM simplifies to a simple linear model; therefore, general purpose solutions to normal linear models are available. This means that the simple LMMs we explored in this paper can easily be generalized to more complex methods used in other GWAS-type applications that currently cannot easily be extended to experiments with heterogeneous samples (e.g. set-tests and multi-marker regressions).

We demonstrated `Grid-LMM` using three examples that are broadly representative of many experimental settings in quantitative genetics. The first was a study of gene-environment interactions, while the second and third focused on the partitioning of genetic variance among additive and non-additive components. Recent studies have shown how neglecting gene-environment covariance when estimating heritability [8] or testing genetic associations [34] can lead to biased estimates. There is little reason to expect these results to be limited to controlled environmental settings, as we have studied here. Previous work has found that incorporating spatial covariance through a Gaussian radial basis function improved estimates of trait heritability within a sample of individuals from Uganda [8]. Spatial variability also exists in agricultural field trials, and two-step approaches are frequently used where spatial variation is removed first and then genetic associations are tested on the residuals. This two-step procedure may be underpowered when true effect sizes are large. Similarly, epistatic and other non-linear genetic variation are known to be large for many traits [59–61] and accounting for this variation may improve our ability to detect both the main effects of markers (as we demonstrated above), as well as possibly interacting loci [17].

The slight differences in posterior means between the `Grid-LMM` and MCMC-based posterior estimates in Fig 3b-c are due to differences in the priors. MCMC-based LMM implementations classically use inverse-Gamma priors for variance components because of conjugacy [20]. However, others have recommended uniform or half-t-family priors for the standard-deviation parameters of hierarchical models [62], which are easily implemented in `Stan` [52]. We used a half-Student-t(3,0,10) distribution for each variance component in our rstan model to produce Fig 3b-c. This is easy to approximate in `Grid-LMM;` relative prior weights can simply be applied to each grid-vertex, resulting in much closer agreement of posterior summaries between the two methods (see S7 Fig). As we show in S8 Fig, supposedly “uniformative” versions of both the inverse-Gamma and half-Cauchy-type priors are actually highly informative for variance component *proportions*. In our experience, it is more natural to elicit priors on variance component proportions than variance components themselves, particularly when the phenotypes are on very different scales, because these can be interpreted as the relative importance of the various factors. This is an additional advantage of the LMM parameterization that we utilize in `Grid-LMM`.

The `Grid-LMM` approach does have limitations. First, the size of the grid spanning the variance components increases nearly exponentially with the number of random effects. Since each grid vertex requires a separate Cholesky decomposition of the observation-level covariance matrix, a grid search quickly becomes prohibitively expensive with more than ~6-8 variance components. This is a general problem for mixed model algorithms, and it may be possible to adapt efficient derivative-based algorithms to the grid space. Our fast-heuristic search method converges on the correct answer in most cases we have examined; however, likelihood surfaces of linear mixed models are not always convex and this algorithm may converge onto a local maximum in such cases. We note that most general-purpose algorithms for LMMs with multiple random effects are also sensitive to this issue. Second, for REML or posterior inference of variance component proportions, `Grid-LMM` estimates are accurate but not precise; specifically, they are limited by the resolution of the grid. We show that this has little impact on hypothesis testing for fixed effects, for example in GWAS settings. However, boundaries of posterior intervals in particular may not be reliable. Nevertheless, summaries like the posterior mean or estimates of the joint posterior density are highly accurate (e.g. Fig 3). Third, the `Grid-LMM` approach is limited to Gaussian linear mixed models. Generalized linear mixed model algorithms rely on iteratively re-weighting the observations, a function that changes the covariance matrix in a way that cannot be discretized. Finally, we have not explored LMMs with correlated random effects, although these are commonly used in quantitative genetics. Since correlation parameters are restricted to the interval (−1, 1), discretizing correlations in parallel with the variance component proportions may be feasible and is an avenue that is worth future study.

## Methods

### Linear Mixed Models

We consider the following parameterization of the standard linear mixed model:

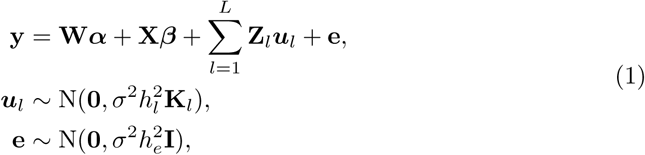

where *n* is the number of observations, *L* is the number of random effect terms (not including the residuals), **y** is an *n ×* 1 vector of quantitative traits, and **W** and **X** are *n × c* and *n × p* design matrices for covariates and marker effects, respectively, with ***α*** and ***β*** corresponding *c ×* 1 and *p ×* 1 vectors of coefficients. Similarly, **Z***_l_* are *n × r_l_* design matrices with corresponding random effects ***u**_l_*, which are normally distributed around zero and have covariance matrices proportional to the known positive semi-definite *r_l_ × r_l_* matrices **K**_*l*_. Lastly, **e** is a *n* × 1 vector of uncorrelated normally distributed errors, and N(•,•) denotes the multivariate normal distribution. The common variance of all random effects are denoted by *σ*^2^, and the vector 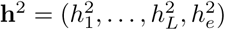 represents the proportion of variance attributed to each random effect term or the residual error. Elements of **h**^2^ are all non-negative and sum to one, forming an *L*-dimensional simplex.

In GWAS applications, we assume that **W** is constant across markers and the value of ***α*** is not of central interest; meanwhile, **X** varies for each test and we aim to perform statistical inference on a subset of ***β***. In heritability estimation applications, we focus on inferring the vector **h**^2^. In both cases, the vectors ***u**_l_* and **e** are nuisance parameters and we can integrate them out resulting in the following equivalent model:

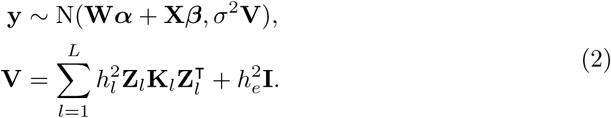

If the matrix **V** is full-rank (which is guaranteed if 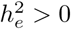), we can use the inverse of the Cholesky decomposition **V** = **LL**^T^ to transform Equation 2 to the following:

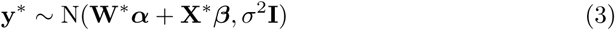

where **y**^∗^= **L**^−1^**y**, **W***^∗^* = **L**^−1^**W** and **X**^*∗*^= **L**^−1^**X**. Equation 3 is a simple linear model for **y**^*∗*^, with the likelihood function:

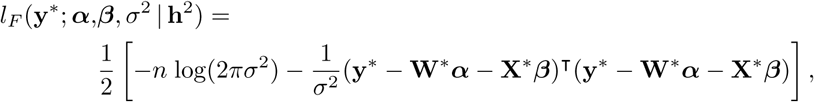

where efficient methods for inference of [***α**, **β***] and *σ*^2^ are well known. We derive maximum-likelihood and restricted-maximum likelihood solutions for these parameters here, as well as posterior distributions under the conditional normal-inverse-gamma prior below. The log-likelihood and restricted-likelihood functions (respectively) for Equation 2 are:

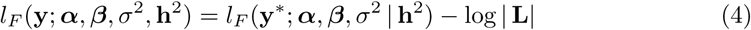

and

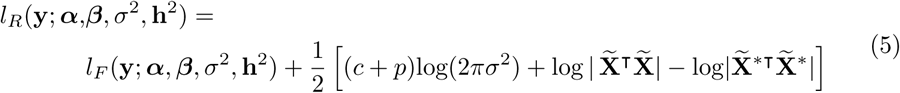

which are simple (and computationally inexpensive) updates to the likelihood function of Equation 3. Here, we denote | • | as the matrix determinant and let 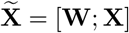 and 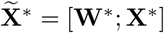, respectively. For ML and REML applications, we can calculate the profile likelihoods *l_F_* (**y**; **h**^2^) and *l_R_*(**y**; **h**^2^) as functions of the profile likelihood of the rotated data, which is formed by maximizing *l_F_* (**y**^*∗*^; ***α**, **β**, σ*^2^ **h**^2^) with respect to ***α***, ***β***, and *σ*^2^. Namely:

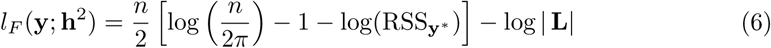

where 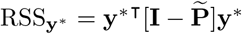 is the residual sum of squares, and 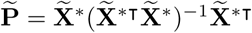 is a projection (hat) matrix.

### Statistical inference by parallel grid search

We now outline the `Grid-LMM` approach for calculating approximate Wald-test statistics, and then show extensions for computing Bayesian posterior distributions of variance components and marker specific Bayes Factors.

A Wald test for the null hypothesis **M*θ***= **0** for ***θ*** = [***α***^T^*, **β***^T^]^T^ and an arbitrary *q ×* (*c* + *p*) matrix **M** uses the general F-statistic:

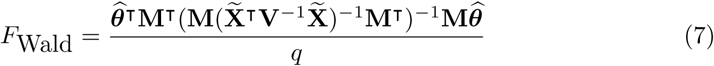

with *q* and (*n-c-p*) degrees of freedom, where 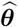 is the estimate of ***θ*** using the REML estimate of **V** [63].

To calculate genome-wide Wald statistics, we must estimate 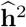 for each of the *p* markers tested. There are no closed-form ML (or REML) solutions for **h**^2^; therefore, iterative algorithms are required. A naive approach involves repeatedly inverting **V**, which scales cubically in the number of observations. Since the estimates for 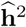 differ for each test, the total computational complexity of in a GWAS setting is *𝒪*(*tpn*^3^) assuming an average of *≈ t* inversions per run of the optimization algorithm (generally ≈ 3-100, increasing with the number of random effect parameters). The fast algorithm used in `GEMMA` [24] and `FaST-LMM` [23] reduces this to *𝒪*(*n*^3^ + *pn*^2^ + *ptc*^2^*n*) by utilizing the eigenvalue decomposition of the similarity matrix **K**. However, this only works for a single random effect. We are unaware of any exact algorithm with lower computational complexity for models with multiple random effects (and full-rank covariance matrices).

With `Grid-LMM`, rather than optimizing **h**^2^ separately for each marker, we instead define a grid of candidate values for **h**^2^ and calculate the restricted profile-likelihood *l_R_*(**h**^2^ **y**^*∗*^) at every grid vertex for each marker. At each grid vertex, we must invert **V** once, but we can re-use this calculation for every marker. This has a computational complexity of approximately *𝒪*(*g*(*n*^3^ + *pn*^2^)) for a grid with *g* vertices. For all analyses reported in the main text, we use a rectangular grid with a resolution of 0.1 or 0.01 *h*^2^-units, with the restriction that all 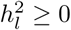 and 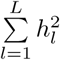. As described below, we either peg this grid to the origin (i.e. 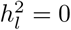, *∀l*), or to the REML estimate 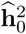 derived from the null model with no marker effects. This grid search generates a vector of *g* profiled Restricted Likelihood scores for each marker. We select the values of 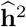 that correspond to the highest such score for each marker and use Equation 7 to calculate the approximate Wald statistics.

To calculate approximate likelihood ratio test statistics, we use the `Grid-LMM` approach to calculate the full profile-likelihoods for models under both the null and alternative hypothesis.

For Bayesian inference, rather than working with the profile likelihoods, we instead use a conditional normal-inverse-gamma prior ([***α***^T^*, **β***^T^]^T^*, σ*^2^) ~ NIG(**0**, **Ψ**, *a*_0_, *b*_0_), and then integrate over the prior to calculate the marginal likelihood of the data given **h**^2^. This results in the following:

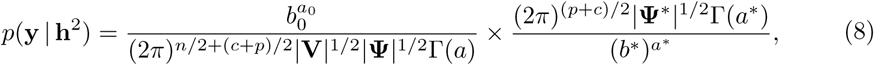

where Γ(•) is the gamma function, 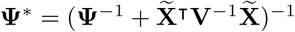, *a^∗^* = *a*_0_ + *n/*2, and *b^∗^* = *b*_0_ + RSS_**y**_*∗,***Ψ***/*2 with RSS_**y**_*∗,***Ψ** having the same form as RSS_**y**_*∗* in Equation 6 — except with 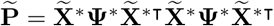. See S1 Supplementary Methods for more detail on the specific derivations. We calculate the marginal likelihood *p*(**y ~ h**^2^) at each vertex of the grid as described above. Assuming a discrete-valued prior *p*(**h**^2^), we can then compute the posterior distribution of **h**^2^ as:

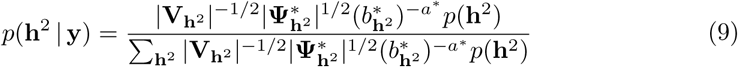

where, for clarity, parameters that are a function of **h**^2^ are denoted with a subscript. Continuous target densities *π*(**h**^2^) can be approximated as *p*(**h**^2^) by assigning each grid vertex a probability equal to the integral of *π*(**h**^2^) over the *L*-dimensional rectangle centered at the corresponding value **h**^2^. We assume a uniform prior over our grid for all analyses presented in the main text. Bayes factors are computed by comparing models under the alternative and null hypotheses as the ratios in Equation 9. All analytical calculations — including the summation in Equation 9 — can be performed on the log-scale to prevent numerical underflows. Terms common to both models drop out of the ratio; therefore, limiting improper priors on ***α*** and *σ*^2^ can be used, which results in scale-independence [49].

### Accelerated grid searches

The full grid search described above is dramatically faster than the naive algorithm for mixed-model GWAS analyses, as long as the vertices of the grid is less than the number of genetic markers (i.e. *g < p*) and can easily be parallelized across multiple computers. However, *g* grows rapidly as the grid resolution and number of random effects increases:

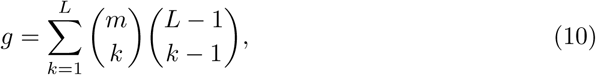

for a grid with *m* divisions per *h*^2^ and, therefore, can still be slow. If we make two assumptions which are commonly true, we can develop heuristics for both the ML/REML and Bayesian algorithms that prevent the need to evaluate every grid vertex:

- The vast majority of markers have little association with the phenotype. This is a common hypothesis in GWAS settings [64–67], and for our purposes, we merely require that the percentage of variance explained individually by most markers is smaller than the difference between grid vertices *≈* 1*/m*.
- The likelihood and/or posterior function is convex. This is not always true, since both the likelihood and posterior functions can have *>* 1 maximum. However, the conditions that cause these events are rare [68], and most exact LMM algorithms are also susceptible to converging to local maxima.

To search for the ML or REML solutions, we first find 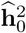 under the null model with no marker effects. We calculate the profile (restricted)-likelihood scores for each test at 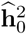, and then form a grid centered at this value by adding or subtracting 1*/m* to each 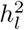 in all combinations. For two random effects, this grid will be a ring around 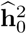 with *g*_1_ *≤* 8 vertices (depending on if 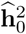 is within 1*/m* of a boundary of the simplex). We calculate the scores for each test at each vertex of the grid, and then compare the maximum scores to the scores at 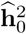. For every test, when no greater value is found, we choose 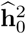 as the maximum and skip the corresponding marker in all future calculations. For the the scores for 1+ tests, and form a new grid of size *g*_2_ around all of them, dropping remaining *p*_2_ markers, we select the set 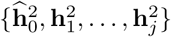 of grid vertices that maximized the scores for 1+ tests, and form a new grid of size *g*_2_ around all of them, dropping vertices already tested. This procedure is repeated *t*_7_ times until the new grid no-longer increases scores for any test and we are confident that all (approximate) maximums have been found. This accelerated search has total complexity 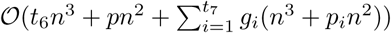, with *t*_6_ the number of iterations needed to optimize variance components under the null model, and *p*_1_ = *p* (see Table 1).

Analogously, to accelerate evaluations of posterior distributions, we evaluate *p*(**y | h**^2^) over an initial grid of resolution 1*/m*_1_ with discrete prior *p_m_* (**h**^2^) and estimate the posterior distribution as in Equation 9. Summary statistics, such as the posterior mean and variance, will be more accurate if the posterior mass is distributed across multiple vertices of the grid. Therefore, we identify the set 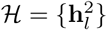 of vertices that together explain 99% of the posterior mass. If the size of this set is below a threshold (say 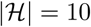), we then form a new grid with double the resolution *m*_2_ = 2*m*_1_ and a new prior *p_m_* (**h**^2^). Vertices that overlap between the grids can be filled in directly. We then begin filling in the new grid by evaluating vertices within 1*/m*_2_ distance from any 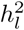 corresponding to the vertices in 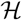. After each iteration, we re-calculate the set 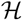, and continue evaluating the neighboring vertices as long as 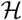 continues to grow. If the size of 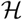 remains smaller than the threshold after whole grid is evaluated, or no new vertices are added to 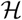 at the end of an iteration, we double the resolution again: *m_i_*_+1_ = 2*m_i_* and repeat the grid-filling steps again. We note that this procedure is only appropriate if the posterior is convex and, therefore, is limited to the case of uniform priors on **h**^2^. A similar procedure was proposed for Bayesian inference in Gaussian process models in the `GPstuff` toolbox, although it is not optimized for parallel inference in GWAS [55].

To accelerate evaluations of GWAS Bayes Factors, we combine the two previously described algorithms. In particular, we define a grid centered on 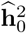 with a resolution 1*/m* that is sufficiently fine such that we expect each posterior to be distributed across multiple vertices. We then calculate *p*(**y | h**^2^) for each test, starting from 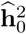 and moving out in concentric rings on the grid. After each iteration, if the new ring contributes *<* 0.01% to the total posterior mass (among evaluated vertices) for that test, we assume the posterior is well characterized and stop evaluating *p*(**y h**^2^) for that marker. As for the ML and REML solutions above, markers with little to no association with the phenotype will lead to posteriors of **h**^2^ that are concentrated close to 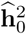; hence, only markers with large effects will shift *p*(**h**^2^ | **y**) strongly to new regions of the grid.

Unless otherwise specified, we used accelerated grid searches for all `Grid-LMM` analyses presented here with grid step sizes of 0.01 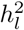 units.

### GWAS datasets

Genotype and phenotype data on 107 *Arabidopsis thaliana* traits and 216,130 genetic markers were downloaded from https://github.com/Gregor-Mendel-Institute/atpolydb/wiki. We follow practices suggested by the original authors of these data and log-transformed a subset of the phenotypes prior to analyses (except for traits that had values less-than or equal to zero) [44]. For the analysis of gene-environment interactions in flowering time, we (1) extracted the trait identifiers “7_FT10” (i.e. growth chamber at 10C) and “57_FT Field” (i.e. field setting), (2) selected data from the 175 accessions that were measured in both environments, and (3) concatenated the two datasets into a single vector of 350 observations. The two traits were individually standardized to have mean zero and standard deviation one prior to analysis. We used the sommer package in R to calculate an additive relationship matrix from all 216,130 markers [69], and then created a G × E kinship matrix as **DZKZ**^T^**D** where **K** is the 175 × 175 additive genomic relationship matrix, **Z** is a 350 × 175 incidence matrix linking observations to accessions, and **D** is a 350 × 350 diagonal matrix with elements equal to −1 or 1 corresponding to observations measured under “7_FT10” and “57_FT Field”, respectively. Plasticities for each accession were calculated as the difference between “57_FT Field” and “7_FT10”. We ran `GEMMA` (version 0.97) with the “-lmm1” option for Wald tests using the **K** matrix described above. We emulated `EMMAX` and `pylmm` functionality by estimating variance components using a null model with no marker effects, and either a single **K** matrix (for single-trait analyses) or 3 random effects (for G × E analysis). Wald test statistics were computed for each marker using our GridLMM R function with a grid consisting of only a single vertex. For G × E analyses, we fit a model with a main effect of the environment, a main effect for each marker of interest, and an interaction between the marker and the environment — and then calculated the Wald-F statistic only for the interaction effect.

The heterogeneous stock of mice data from the Wellcome Trust Centre for Human Genetics (http://mtweb.cs.ucl.ac.uk/mus/www/mouse/index.shtml) consists of 1,814 individuals from 85 families, all descending from eight inbred progenitor strains [46]. We used the marker and phenotype data provided in the `BGLR R` package [70] from which we extracted the “EndNormalBW” trait and information on the gender and cage of each mouse. Gender was used as a covariate in all analyses, and cage assignment was treated as a random effect in the three-random-effect models. Additive and epistatic kinship matrices were calculated using `sommer`. Wald test statistics and Bayes Factors were calculated using the heuristic (accelerated) grid search of `Grid-LMM`. Bayes Factors were calculated assuming flat priors on the intercept and residual variance term, a standard normal prior N(0, 1) on the marker effects — similar to the previously proposed *D*_2_ prior [49] — as well as a uniform prior over the grid of variance component proportions.

Computational timings for GWAS analyses are reported for a MacBookPro14,3 with a 2.9Ghz Intel Core i7 processor and using only a single CPU core. Further speedups are possible by parallelizing the grid search. For accelerated grid searches, REML estimates of variance components under the null model (i.e. the starting points of the grid search) were calculated with LDAK and are included in the computational time estimates.

### Gene expression dataset

Gene expression data on 24,175 genes from 728 *Arabidopsis* accessions was downloaded from http://signal.salk.edu/1001.php and subsetted to genes with average counts ≥10. A genomic relationship matrix (**K**_*A*_) for 1,135 accessions was downloaded from http://1001genomes.org/data/GMI-MPI/releases/v3.1/SNP_matrix_imputed_hdf5/. Both sets of data were subsetted to an overlapping set of 665 accessions. **K**_*A*_ was then centered by projecting out the intercept and normalized to have mean diagonal elements equal to one. The pairwise-epistasis genomic relationship matrix **K**_*E*_ was calculated with element-wise multiplication as **K**_*A*_ ⊙ **K**_*A*_, and then also normalized to have mean diagonal elements equal to one. The gene expression matrix was normalized and variance-stabilized using the `varianceStabilizingTransformation` function of the `DEseq2` package [71]. Grid-LMM REML estimates were compared to exact REML estimates from the `LDAK` program with variance components constrained to be non-negative. Grid-LMM posterior distributions estimates were compared to those estimated using `Stan` [52] with the `rstan R` package [53]. To speed computation, we diagonalized the **K**_*A*_ covariance matrix by calculating the singular value decomposition **K**_*A*_ = **USU**^T^, and pre-multiplying both sides of the LMM by **U**^T^. We used a half-Student-t prior distribution with 3 degrees of freedom and a scale of 10 for the three standard deviation parameters. We ran four MCMC chains each of length 10,000, with the first 5,000 as warmup and a thinning rate of 20. Because of poor mixing, this resulted in an “n_eff_” of approximately 200-400 per variance component parameter for each trait.

### Power simulations

We compared the power of one and two-random effect models for detecting G × E markers based on the Arabidopsis flowering time data. We calculated additive genetic and G × E relationship matrices as described above using the actual genotypes. We then simulated phenotypes by summing together a G × E effect of a particular marker, (co)variance from the two random effects, and normally distributed error. For each combination of marker effect sizes ({0, 0.025, 0.05, 0.1, 0.15, 0.2} % of the total phenotypic variance), and random effect proportions (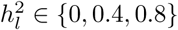 for each random effect), we ran 10,000 simulations with different randomly selected markers from the Arabidopsis genotype data. We then fit five models to each dataset and calculated Wald-tests for marker G × E effects using each. The five methods consisted of three two-random effect approaches including: (1) an exact two-random effect model fit with `LDAK`, (2) the approximate two-random effect model fit with `Grid-LMM` with a grid size of 0.1 *h*^2^-units, and (3) the approximate two-random effect model `pylmm` that conditions on variance components estimated under the null model [34]. We also consider two one-random effect models that could be fit with `GEMMA`: (4) a model that only included the additive genetic relationships and ignored the G × E covariance, and (5) a model that only included the G × E covariance and ignored the additive genetic relationships.

For each method and for each simulation scenario, we first calculated a genomic control inflation factor [1] for the GxE marker tests as the ratio between the median value of the the *F*-statistics returned by each model and the median value of a *F*_1,312_ distribution since *n* = 316 and each model included *p* = 4 predictors (overall intercept, environmental effect, the main effect of the marker, and the G × E interaction between the environment and the marker). We then “corrected” all *F*-statistics by dividing by the appropriate inflation factor, and calculated statistical power for each GWAS method as the proportion of corrected Wald test p-values exceeding the genome-wide significance level at the conventional Bonferroni corrected threshold *P* = 2 × 10^−7^ (see S4 Fig).

The results show that `Grid-LMM` maintains nearly identical power to the exact method in each simulation, all while also maintaining well-calibrated p-values under the null-distribution. The approximate method based on `pylmm` has uniformly lower power, but maintains an accurate null-distribution. The one-random effect methods show opposite results: when including only the additive relationship matrix, the null-distribution of *−*log10(*p*)-values is greatly inflated when G*×*E-variation is large. On the other hand, when only G*×*E variance is modeled and yet the additive genetic variance is large, −log10(*p*)-values are greatly deflated. These results all confirm our `expectations` that modeling covariance accurately, but not necessarily precisely, is important for accurate association mapping in GWAS.

### Sample size simulations

We developed a simulation strategy to evaluate the effect of sample size on the accuracy of `Grid-LMM` in a reasonable amount of time without the confounding issue of changes in population structure across different populations. These simulations are based on the WTCHG body weight data described above, creating simulations with similar levels of structure and random effect covariance as in the real analysis, and comparing the accuracy of marker tests using the `Grid-LMM` and `null-LMM` methods to the `Exact-LMM method`.

We began with our largest dataset (i.e. the *n* = 1814 WTCHG heterogeneous stock mice), randomly selected 300 markers, and calculated the three covariance matrices for additive, addtive-additive epistasis, and cage random effects. Next, we created a larger dataset of 5 × the original size by repeating the original marker data five times and similarly created three block-diagonal covariance matrices by repeating each original covariance matrix five times. While not completely realistic, this created a population similar to what we would expect if the heterogeneous stock population was created five separate times from five independent sets of progenitors; therefore, it has a similar level of structure relative to its size as the original *n* = 1, 814 population. Finally, we created a smaller dataset of 1*/*5× the size by subsampling the first 362 individuals from this dataset, their corresponding marker data, and the corresponding partitioned subsets of the three covariance matrices.

For each simulation, we selected a single marker and assigned it an effect size between 0 and 0.15 in 16 steps. We then added four random vectors corresponding to the three random effects and “iid” error. To be realistic, we used the variance component proportions 0.23, 0.29, and 0.25, respectively, as weights for the random vectors (with the sum scaled so that the total phenotypic variance equaled one). This choice was based on the observed variance components in the real bodyweight data. We repeated this simulation strategy for each of the 300 markers in the three populations and each of the 16 effect sizes. Within each simulation, we calculated marker p-values using the Exact-LMM and null-LMM methods. We then simulated Grid-LMM results by perturbing each of the three variance component proportions from Exact-LMM: ± 0.05 for Grid-LMM(0.1) and ±0.005 Grid-LMM(0.01), respectively. Lastly, we selected the p-value from the model with the highest REML score. This represented a “worst-case scenario” for Grid-LMM where the optimal variance components were maximally far from the grid vertices.

To estimate the time for a GWAS under each population size, we measured the length of time to fit `Exact-LMM` for a single marker using LDAK (*T*_1_), the time to perform a single Cholesky decomposition (*T*_2_), and the time to calculate p-values for a set of *p* markers given a pre-calculated Cholesky decomposition *pT*_3_. We then calculated the total time for a GWAS with *p* markers as:

1. `Exact-LMM`: *pT*_1_
2. `null-LMM`: *T*_1_ + *T*_2_ + *pT*_3_
3. `Grid-LMM`(0.1): *g*_1_(*T*_2_ + *pT*_3_)
4. `Grid-LMM-Fast`(0.01): *T*_1_ + *g*_2_(*T*_2_ + *pT*_3_) + *g*_3_(*T*_2_ + *T*_3_)

where *g*_1_ = 220 is the size of a complete grid for three random effects with resolution *h*^2^ = 0.1, *g*_2_ = 27 is the size of a ball around the `null-LMM` variance component estimates in the `Grid-LMM-fast` heuristic, and *g*_3_ is the number of additional grid vertices that must be traversed by the `Grid-LMM-fast` algorithm. For the `Grid-LMM-fast` calculations, we assumed that only a single marker had a non-zero effect (and so only this marker would need to be taken through multiple iterations of the heuristic search), and that the effect size of this marker was approximately at the power threshold given the sample size (≈ 0.1 for *n* = 362, ≈ 0.035 for *n* = 1814, ≈ 0.006 for *n* = 9070; see Fig 4b).

## Software Availability

Software for computing the Grid-LMM is carried out in R code, which is freely available at https://github.com/deruncie/GridLMM. Scripts for running the analyses reported in the manuscript are available at https://github.com/deruncie/GridLMM_scripts.

## Author Contributions Statement

D.E.R. and L.C. conceived the experiment(s), D.E.R. conducted the experiment(s), D.E.R. and L.C. analyzed the results. Both authors wrote and revised the manuscript.

## Supporting information

**S1 Fig. Approximate time and memory requirements of** `Grid-LMM` **as a function of sample size.** (**a**) Computational times for the most costly steps of typical mixed model fitting algorithms: inverting an *n × n* covariance matrix (generally using a Cholesky decomposition), and multiplying the inverse matrix by an *n*-vector (i.e. a vector of marker genotypes). The red curve shows the time required for a Cholesky decomposition using the base R function chol as a function of *n*. The green curve shows the time required for a Cholesky matrix-by-marker matrix multiplication with 1 × 10^5^ markers, as a function of *n*. The blue curve is the sum of the Cholesky decomposition and matrix-vector multiplication operations for a single grid-cell in `Grid-LMM` with 1 × 10^5^ markers. The green and blue curves are barely distinguishable across most of the range because the Cholesky decomposition is generally not limiting. The purple curve would be the expected time for a separate Cholesky decomposition and matrix-vector multiplication for *each marker* in a GWAS with 1 × 10^5^ markers (i.e. the cost of a typical `exact-LMM` method such as `LDAK` for a single iteration). Both the `Grid-LMM` and LDAK times are *per-iteration*. `Grid-LMM` requires this time at each grid cell. `LDAK` requires multiple iterations for the REML optimization separately for each marker. Generally, `Grid-LMM` will evaluate more grid cells than `LDAK` requires iterations per test. However this will not cause a reversal in the relative time requirements unless a very large grid is used. (**b**) Memory requirements for storing an *n × n* Cholesky matrix as a function of sample size. In both panels, the curves were extrapolated based on tests with *n* between 256 and 4096 (actual times shown with points). All timings were estimated using base R functions.

**S2 Fig. Accuracy of the log-transformed *p*-values across the 107 Arabidopsis phenotypes [44].** GWASs were run for each phenotype using 216,130 markers and up to 199 accessions, with a single random effect controlling for additive genetic relationships among lines. For each phenotype (represented as a single point in the plots), we compared the exact Wald-test −log_10_(*p*) calculated by `GEMMA` to p-values calculated by the approximate methods `EMMAX`, and `Grid-LMM` using either the naive approach with a complete grid of size 0.1 *h*^2^-units, or the fast heuristic algorithm `Grid-LMM-fast` with a fine grid size of 0.01-*h*^2^ units. `Grid-LMM` p-values were always at least as accurate (as measured by root mean-squared-error, RMSE) as those calculated by `EMMAX`. Specifically, p-values calculated with a fine grid-size of 0.01 (using the fast algorithm) were nearly indistinguishable from those of `GEMMA`, except in the rare cases where the REML surface was not unimodal. This was generally restricted to a small subset of rare markers with small-moderate effect sizes, and only occurred for a few traits. In these cases, the complete — but more coarse — grid search of `Grid-LMM` with step sizes of 0.1 was more accurate.

**S3 Fig. Genomic control inflation factors for simulated data.** Simulated datasets were created based on the Atwell genotype data and the G × E analysis. We randomly selected 10,000 markers, generated simulated data with different proportions of additive (G) and gene-environment interaction (G × E) variation for each marker, and calculated Wald *F*-statistics for an interaction between the marker and the environment. Bars show an estimate of genomic control inflation factors [1] for each of the following five methods. `exact-LMM-G+GxE` is an exact LMM algorithm fit with `LDAK`. This model included both random effects and the marker effect. At a genome-wide scale, it is very slow, with computational complexity 𝒪(*ptn*^3^). `exact-LMM-G` and `exact-LMM-GxE` are exact LMM algorithms, similar to `GEMMA`, which included only one random effect and the marker effect. `null-LMM` is an approximate method similar to `pylmm` that conditions on variance components estimated under a null model with no marker effect. It was run with both random effects. `Grid-LMM` was run with a grid size of 0.1 *h*^2^-units and included both random effects and the marker effect. The *λ* values were calculated as the ratio between the median value of the the *F*-statistics returned by each model and the median value of a *F*_1,316*−*4_ distribution. The horizontal line shows the expected value *λ* = 1 under the true model.

**S4 Fig. Power analysis for simulated data.** Bars show the genome-wide power for randomly selected SNPs in the Atwell genotype data under simulations with different proportions of additive (G) and gene-environment interaction (G × E) variation with different marker effect sizes. Simulations were generated as described in S3 Fig, and included only a single marker with zero main effect and G × E effects scaled to a defined percentage of the phenotypic variation. The remaining phenotypic variation was simulated from a multivariate normal distribution constructed by appropriately weighting the additive relationship matrix, the G × E covariance matrix, and the uncorrelated residual variation. Each simulation was run separately for 10,000 randomly selected markers. Wald *F*-statistics from each method were normalized by dividing by the genomic control inflation factor computed for Figure, and then *p-values* were calculated and compared to the Bonferroni corrected threshold *P* = 2 × 10^−7^ to determine significance.

**S5 Fig. Comparison of REML estimates between** `Grid-LMM` **and** `LDAK` **for 20,843 *Arabidopsis* genes.** (**a**) REML estimates for the additive genetic variance (variance component for **K**_*A*_). (**b**) REML estimates for the epistatic genetic variance (variance component for **K**_*E*_).

**S6 Fig. Comparison of REML estimates for the additive genetic variance between models with and without an additional pairwise-epistasis random effect for 20,843 *Arabidopsis* genes.** Both models were fit using `Grid-LMM` with a grid size of 0.05 *h*^2^ units. Point positions are jittered for clarity.

**S7 Fig. Posterior distributions of variance components for two genes under the half-Student-t(3,0,10) prior.** Panels (**b**) and (**c**) of Figure 3 in the main text are repeated, except the half-Student-t(3,0,10) prior on the standard deviation of the variance components of **K**_*A*_ and **K**_*E*_ for the random effects was applied to each grid vertex. The prior was approximated by simulating 1 *×* 10^4^ independent draws for *σ_A_*, *σ_E_* and *σ_e_*, converting these to prior draws for 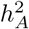 and 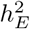, and then measuring the proportion of draws closest to each grid vertex.

**S8 Fig. Inverse-Gamma and half-Student-t priors are informative for variance component proportions.** We compare the implied prior distributions on variance component proportions for three classes of priors in a two-random effect model (e.g. **K**_*A*_ and **K**_*E*_ as random effects plus uncorrelated random error). (**a**)-(**b**) Each standard deviation parameter was assigned a half-Student-t prior with 3 degrees of freedom and scale parameter of 10. (**c**)-(**d**) Each variance parameter was assigned an inverse-Gamma prior with shape parameter 2 and scale parameter 1. (**e**)-(**g**) A uniform prior was applied to the 2-dimensional simplex of 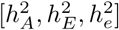. This is the default prior in GridLMM and equivalent to all analyses reported in the main text. (**a**)-(**c**)-(**e**) 2D-density plots for the two variance component proportions. Lighter blue denotes higher prior density. (**b**)-(**d**)-(**f**) Marginal densities for the **K**_*A*_ variance component proportion under each prior. The half-Student-t prior implies high probability that only one variance component is important. The inverse-Gamma prior implies high probability that all variance component proportions are non-zero.

**S1 Table Markers associated with heterogenous stock mice body weight only when accounting for non-additive genetic covariance.** Markers with −log_10_(*p*) *>* 3 on chromosomes 4 and 11 are shown. Position information for each marker is derived from [72]. Other studies that identified the same marker associations are listed. WTC: Wellcome Trust Center Heterogeneous Stock Mice. LG,SM Advanced Intercross: F_9_ and F_10_ generation mice from of an intercross between LG/J and SM/J strains selected for differences in body size.

**S1 Supplementary Methods** Derivation of Bayesian posterior and Bayes Factors.

