## Supplemental figures for "Fast and flexible linear mixed models for genome-wide genetics"

### Grid-LMM: Supplemental Note

#### Figures

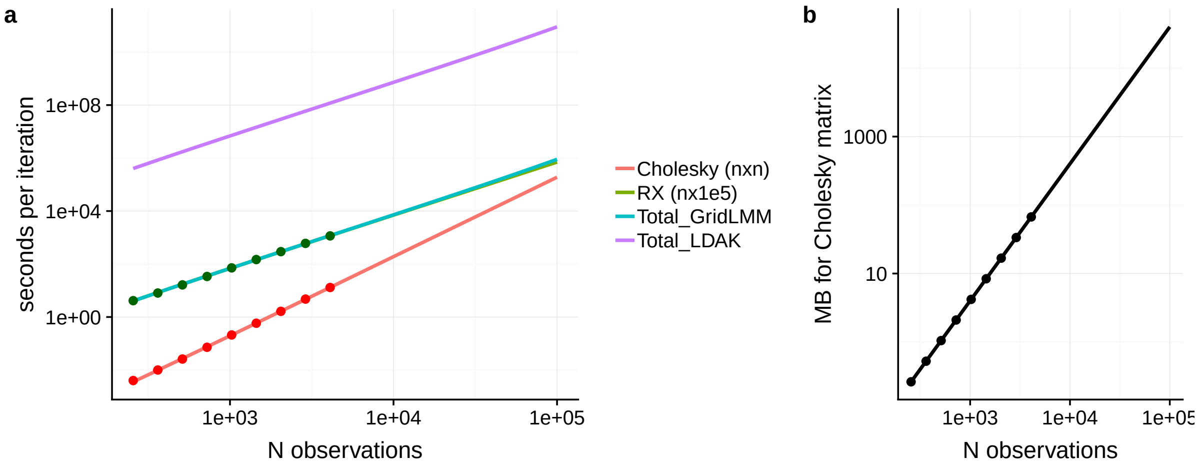


**Figure S1. Approximate time and memory requirements of `Grid-LMM` as a function of sample size.** (**a**) Computational times for the most costly steps of typical mixed model fitting algorithms: inverting an $n \times n$ covariance matrix (generally using a Cholesky decomposition), and multiplying the inverse matrix by an $n$-vector (i.e. a vector of marker genotypes). The red curve shows the time required for a Cholesky decomposition using the base `R` function `chol` as a function of $n$. The green curve shows the time required for a Cholesky matrix-by-marker matrix multiplication with $1 \times 10^5$ markers, as a function of $n$. The blue curve is the sum of the Cholesky decomposition and matrix-vector multiplication operations for a single grid-cell in `Grid-LMM` with $1 \times 10^5$ markers. The green and blue curves are barely distinguishable across most of the range because the Cholesky decomposition is generally not limiting. The purple curve would be the expected time for a separate Cholesky decomposition and matrix-vector multiplication for *each marker* in a GWAS with $1 \times 10^5$ markers (i.e. the cost of a typical `exact-LMM` method such as `LDAK` for a single iteration). Both the `Grid-LMM` and LDAK times are *per-iteration*. `Grid-LMM` requires this time at each grid cell. `LDAK` requires multiple iterations for the REML optimization separately for each marker. Generally, `Grid-LMM` will evaluate more grid cells than `LDAK` requires iterations per test. However this will not cause a reversal in the relative time requirements unless a very large grid is used. (**b**) Memory requirements for storing an $n \times n$ Cholesky matrix as a function of sample size. In both panels, the curves were extrapolated based on tests with $n$ between 256 and 4096 (actual times shown with points). All timings were estimated using base `R` functions.

-
-

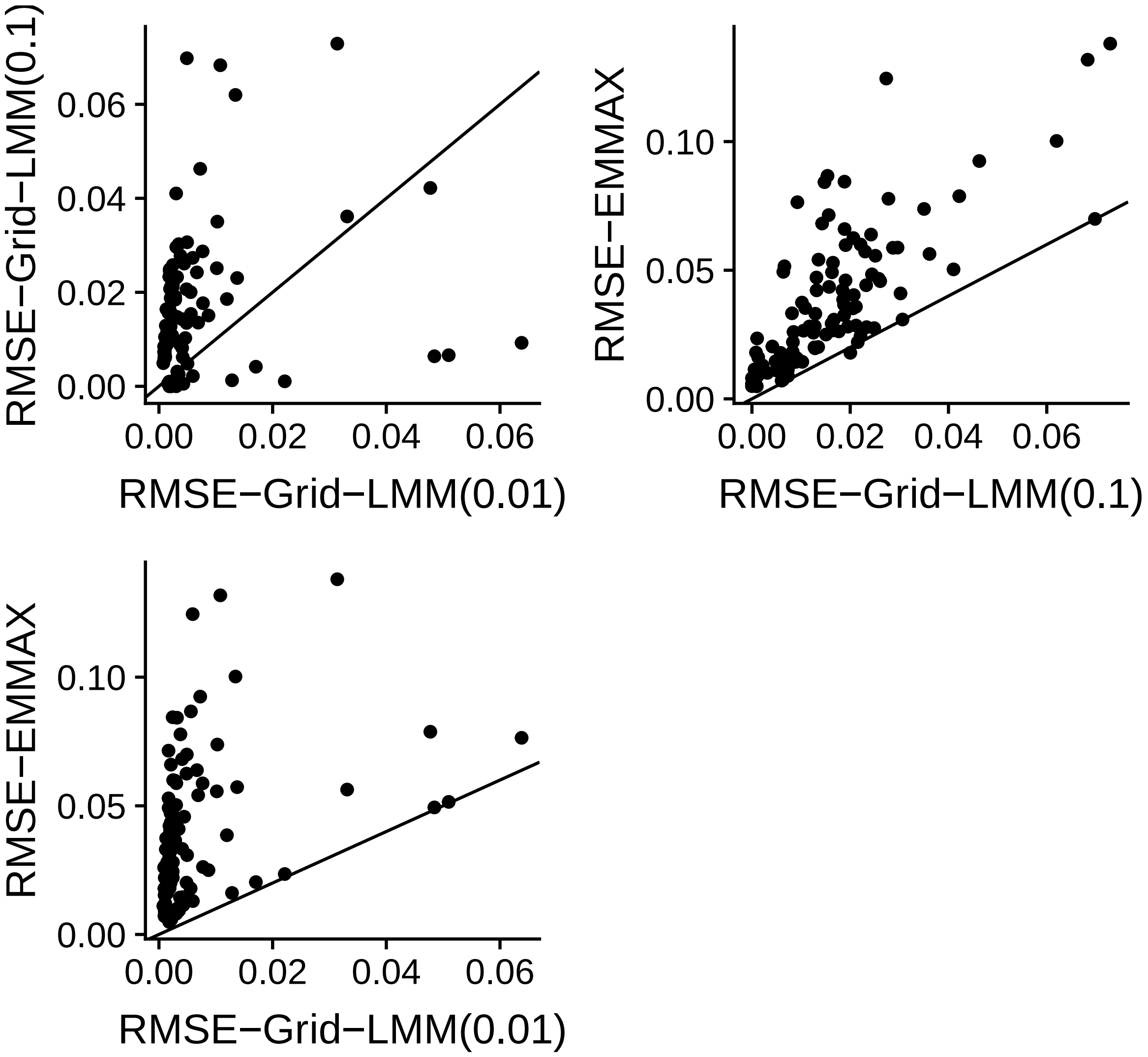


**Figure S2.** **Accuracy of the log-transformed *p*-values across the 107 Arabidopsis phenotypes [1].** GWASs were run for each phenotype using 216,130 markers and up to 199 accessions, with a single random effect controlling for additive genetic relationships among lines. For each phenotype (represented as a single point in the plots), we compared the exact Wald-test -$\log_{10}(p)$ calculated by `GEMMA` to p-values calculated by the approximate methods `EMMAX`, and `Grid-LMM` using either the naive approach with a complete grid of size 0.1 $h^2$-units, or the fast heuristic algorithm `Grid-LMM-fast` with a fine grid size of 0.01-$h^2$ units. `Grid-LMM` p-values were always at least as accurate (as measured by root mean-squared-error, RMSE) as those calculated by `EMMAX`. Specifically, p-values calculated with a fine grid-size of 0.01 (using the fast algorithm) were nearly indistinguishable from those of `GEMMA`, except in the rare cases where the REML surface was not unimodal. This was generally restricted to a small subset of rare markers with small-moderate effect sizes, and only occurred for a few traits. In these cases, the complete — but more coarse — grid search of `Grid-LMM` with step sizes of 0.1 was more accurate.

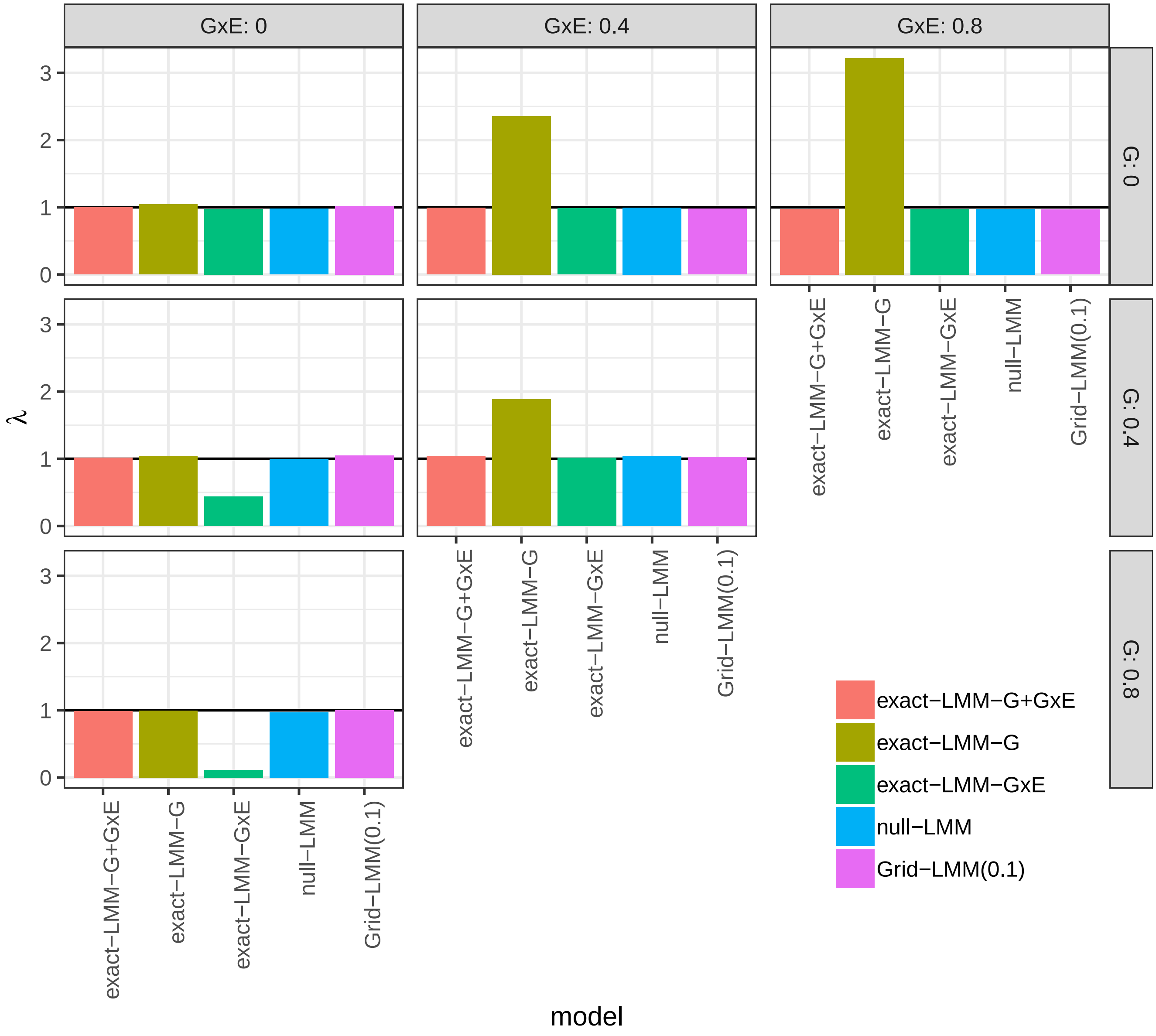


**Figure S3.** **Genomic control inflation factors for simulated data.** Simulated datasets were created based on the Atwell genotype data and the G×E analysis. We randomly selected 10,000 markers, generated simulated data with different proportions of additive (G) and gene-environment interaction (G× E) variation for each marker, and calculated Wald $F$-statistics for an interaction between the marker and the environment. Bars show an estimate of genomic control inflation factors for each of the following five methods. `exact-LMM-G+GxE` is an exact LMM algorithm fit with `LDAK`. This model included both random effects and the marker effect. At a genome-wide scale, it is very slow, with computational complexity $\mathcal{O}(ptn^3)$. `exact-LMM-G` and `exact-LMM-GxE` are exact LMM algorithms, similar to `GEMMA`, which included only one random effect and the marker effect. `null-LMM` is an approximate method similar to `pylmm` that conditions on variance components estimated under a null model with no marker effect. It was run with both random effects. *Grid-LMM* was run with a grid size of 0.1 $h^2$-units and included both random effects and the marker effect. The $\lambda$ values were calculated as the ratio between the median value of the the $F$-statistics returned by each model and the median value of a $F_{1,316-4}$ distribution. The horizontal line shows the expected value $\lambda = 1$ under the true model.

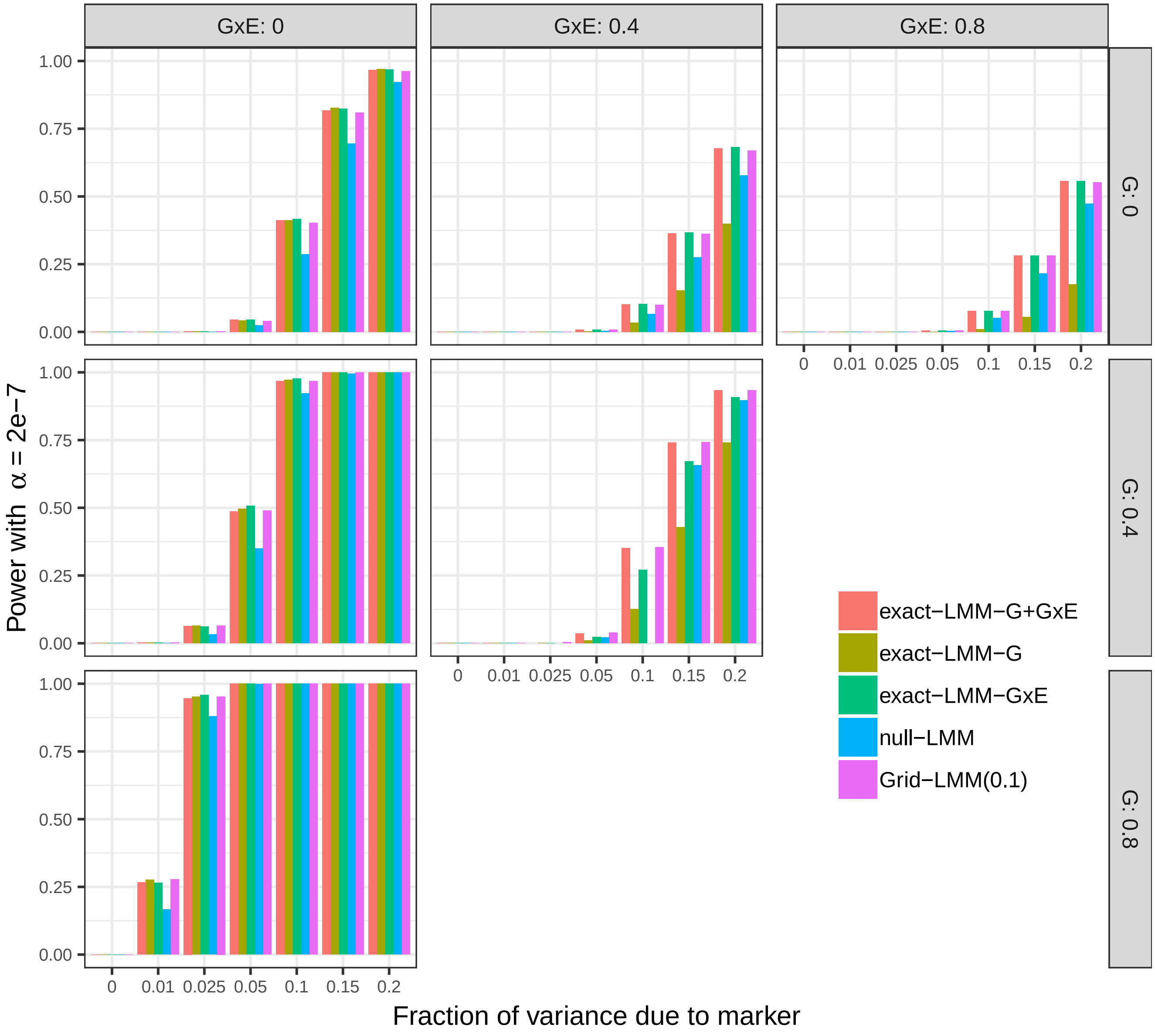


**Figure S4. Power analysis for simulated data.** Bars show the genome-wide power for randomly selected SNPs in the Atwell genotype data under simulations with different proportions of additive (G) and gene-environment interaction (G×E) variation with different marker effect sizes. Simulations were generated as described in Figure S3, and included only a single marker with zero main effect and G×E effects scaled to a defined percentage of the phenotypic variation. The remaining phenotypic variation was simulated from a multivariate normal distribution constructed by appropriately weighting the additive relationship matrix, the G×E covariance matrix, and the uncorrelated residual variation. Each simulation was run separately for 10,000 randomly selected markers. Wald $F$-statistics from each method were normalized by dividing by the genomic control inflation factor computed for Figure S3, and then *p-values* were calculated and compared to the Bonferroni corrected threshold $P = 2 \times 10^{-7}$ to determine significance.

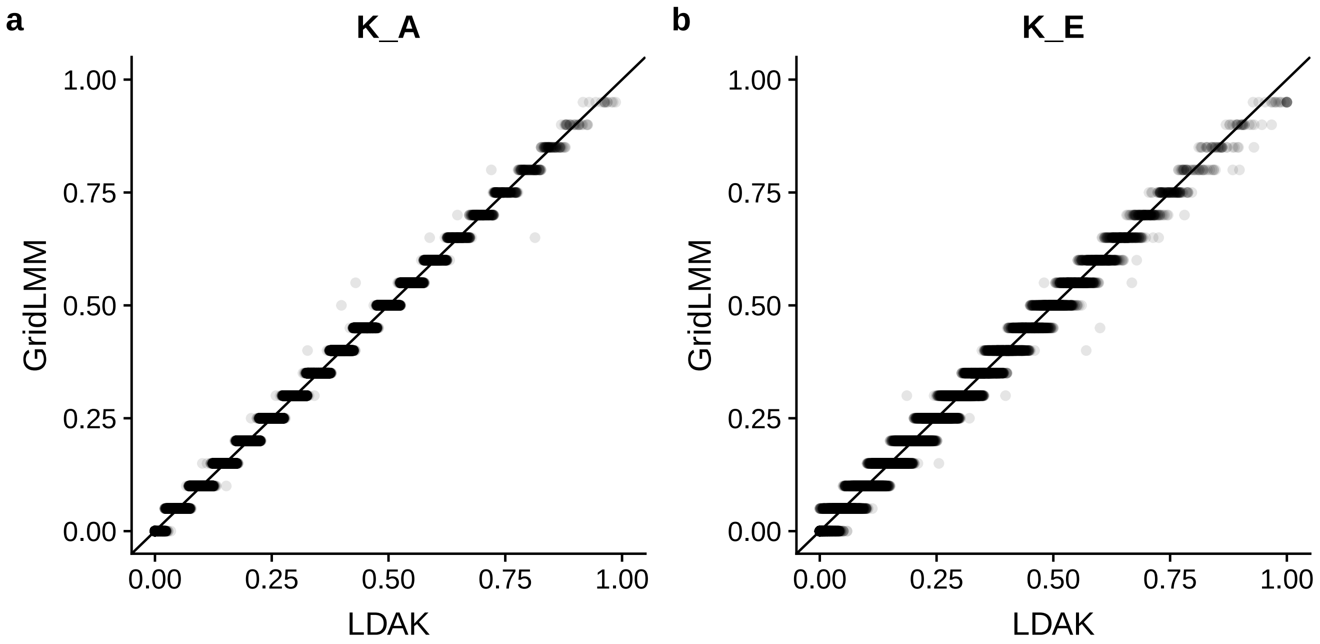


**Figure S5. Comparison of REML estimates between Grid-LMM and LDAK for 20,843 *Arabidopsis* genes.** (**a**) REML estimates for the additive genetic variance (variance component for $\mathbf{K}_A$). (**b**) REML estimates for the epistatic genetic variance (variance component for $\mathbf{K}_E$).

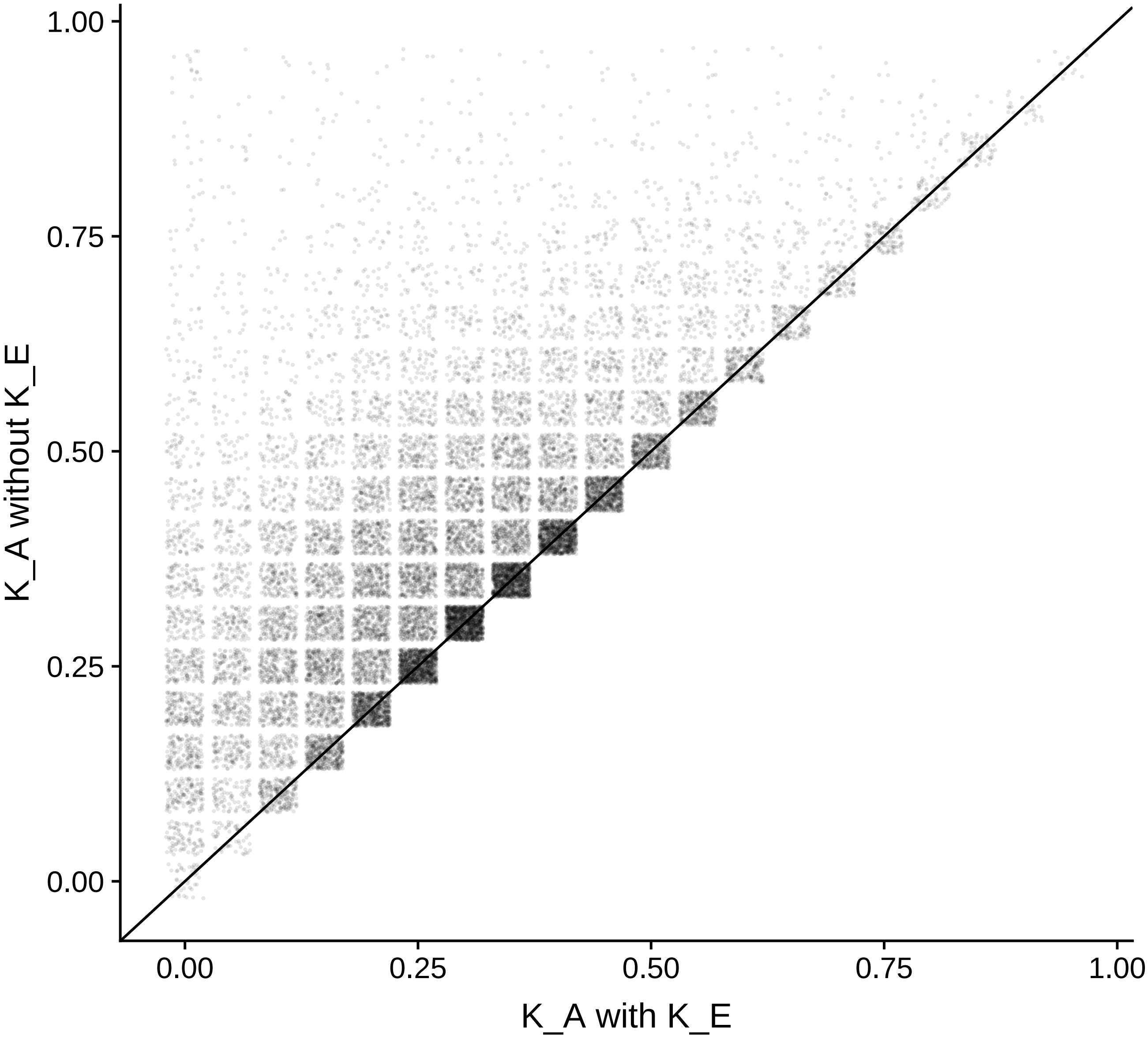


**Figure S6.** **Comparison of REML estimates for the additive genetic variance between models with and without an additional pairwise-epistasis random effect for 20,843 *Arabidopsis* genes.** Both models were fit using Grid-LMM with a grid size of 0.05 $h^2$ units. Point positions are jittered for clarity.

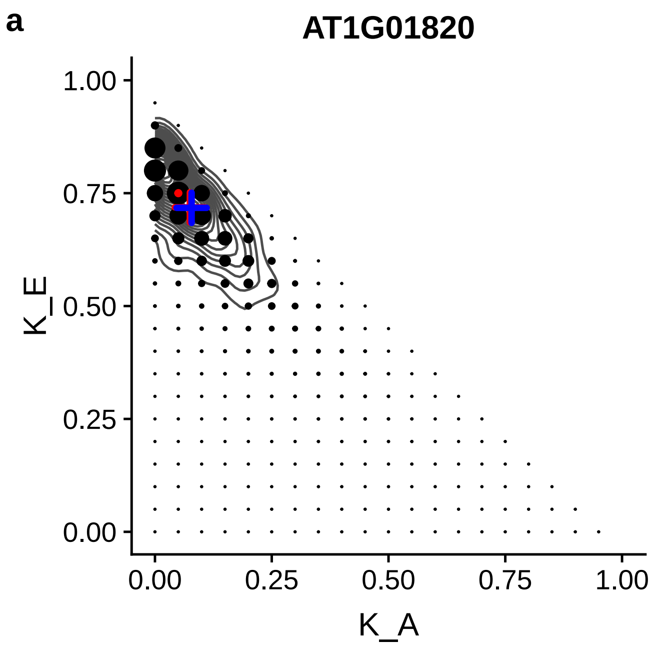


**Figure S7.** **Posterior distributions of variance components for two genes under the half-Student-t(3,0,10) prior.** Panels (**b**) and (**c**) of Figure 3 in the main text are repeated, except the half-Student-t(3,0,10) prior on the standard deviation of the variance components of $\mathbf{K}_A$ and $\mathbf{K}_E$ for the random effects was applied to each grid vertex. The prior was approximated by simulating $1 \times 10^4$ independent draws for $\sigma_A$, $\sigma_E$ and $\sigma_e$, converting these to prior draws for $h^2_A$ and $h^2_E$, and then measuring the proportion of draws closest to each grid vertex.

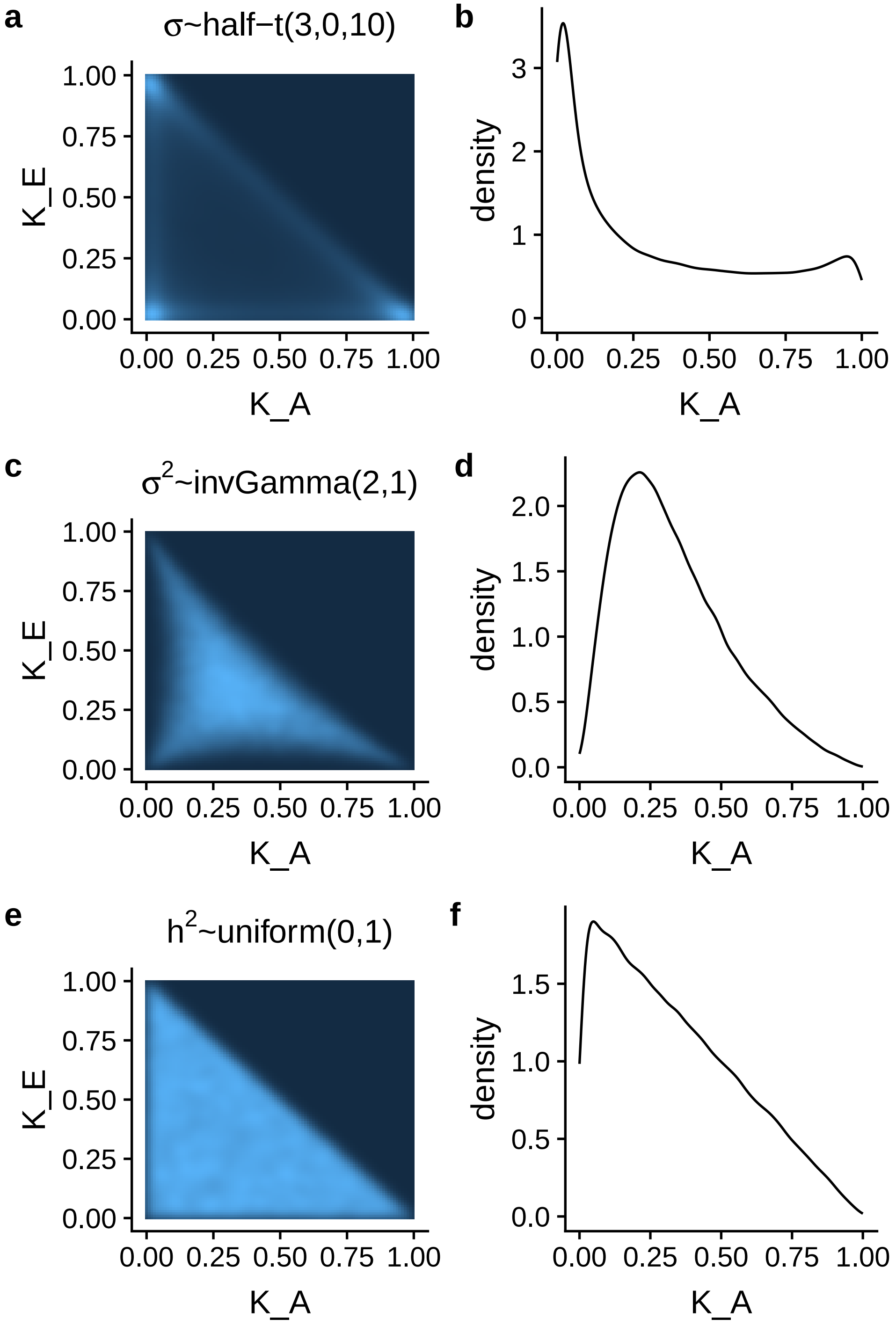


**Figure S8. Inverse-Gamma and half-Student-t priors are informative for variance component proportions.** We compare the implied prior distributions on variance component proportions for three classes of priors in a two-random effect model (e.g. $\mathbf{K}_A$ and $\mathbf{K}_E$ as random effects plus uncorrelated random error). (**a**)-(**b**) Each standard deviation parameter was assigned a half-Student-t prior with 3 degrees of freedom and scale parameter of 10. (**c**)-(**d**) Each variance parameter was assigned an inverse-Gamma prior with shape parameter 2 and scale parameter 1. (**e**)-(**g**) A uniform prior was applied to the 2-dimensional simplex of $[h^2_A, h^2_E, h^2_e]$. This is the default prior in GridLMM and equivalent to all analyses reported in the main text. (**a**)-(**c**)-(**e**) 2D-density plots for the two variance component proportions. Lighter blue denotes higher prior density. (**b**)-(**d**)-(**f**) Marginal densities for the $\mathbf{K}_A$ variance component proportion under each prior. The half-Student-t prior implies high probability that only one variance component is important. The inverse-Gamma prior implies high probability that all variance component proportions are non-zero.

#### Tables

**Table S1. Markers associated with heterogenous stock mice body weight only when accounting for non-additive genetic covariance.** Markers with -$\log_{10}(p) > 3$ on chromosomes 4 and 11 are shown. Position information for each marker is derived from [2]. Other studies that identified the same marker associations are listed. WTCHG: Wellcome Trust Center Heterogeneous Stock Mice. LG,SM Advanced Intercross: $F_9$ and $F_{10}$ generation mice from of an intercross between LG/J and SM/J strains selected for differences in body size.

| marker | Chr | bp | cM | Effect Size | -log10(p) | Genes | References | Phenotype | Population |
|---|---|---|---|---|---|---|---|---|---|
| rs4224463 | 4 | 44561913 | 20.8151 | -0.2307899 | 5.827733 | Trmt10b, Exosc3 | [3] | Starting weight | WTCHG |
| rs6313392 | 4 | 44568868 | 20.81513 | -0.2307899 | 5.827733 | Exosc3 | [3] | Starting weight | WTCHG |
| rs3665393 | 4 | 44721094 | 20.81515 | -0.2313425 | 5.941238 | Shb | [3] | Starting weight | WTCHG |
| rs13477678 | 4 | 44734525 | 20.81518 | -0.1715087 | 3.978578 | Trim14, Coro2a, Shb | — | — | — |
| rs3668228 | 4 | 44764150 | 20.8152 | -0.2307148 | 5.794518 | Shb | [3] | Starting weight | WTCHG |
| rs13477682 | 4 | 45785592 | 21.78908 | -0.1339141 | 3.199823 | Gm16731 | — | — | — |
| rs13481023 | 11 | 51651037 | 29.3193 | 0.1682438 | 4.543509 | Gm26551, Cdkl3 | [4] | Tibia length | LG,SM Advanced Intercross |
| rs8243055 | 11 | 51893791 | 29.31933 | 0.1682438 | 4.543509 | Gm26551, Tcf7 | — | — | — |
| rs6173994 | 11 | 52168577 | 29.51397 | 0.1599403 | 4.19647 | — | — | — | — |
| rs3668680 | 11 | 55712245 | 31.26964 | -0.1584908 | 3.386805 | Gm12239 | — | — | — |
