## Supplemental methods for "Fast and flexible linear mixed models for genome-wide genetics"

### Bayesian posterior and Bayes Factor derivation

We describe our approach for approximating Bayesian posteriors of variance component proportions, as well as calculating Bayes Factors for linear mixed model (LMM) comparisons, using Grid-LMM. Consider the following Bayesian hierarchical regression model specification:

$$\begin{aligned} \mathbf{y} &= \mathbf{X}\boldsymbol{\beta} + \mathbf{e} \\ \mathbf{e} &\sim \mathrm{N}(\mathbf{0}, \sigma^2\mathbf{V}) \\ \boldsymbol{\beta} &\sim \mathrm{N}(\boldsymbol{\mu}, \sigma^2\boldsymbol{\Psi}) \\ \sigma^2 &\sim \mathrm{IG}(a,b), \end{aligned} \tag{1}$$

where $\mathbf{y}$ is an $n\times 1$ vector of observations, $\mathbf{X}$ is an $n\times p$ design matrix, $\boldsymbol{\beta}$ is an $p\times 1$ coefficient vector, and $\mathbf{e}$ is an $n\times 1$ vector of independent and identically distributed errors. The joint prior distribution $p(\boldsymbol{\mu},\sigma^2)$ is said to be the normal-inverse-gamma NIG$(\boldsymbol{\mu},\boldsymbol{\Psi},a,b)$ distribution. The residual covariance matrix $\mathbf{V} = \sum_{l=1}^{L} h_l^2\mathbf{K}_l + h_e^2\mathbf{I}$ is the result of integrating over all random effects, and $h_e^2 + \sum_{l=1}^{L} h_l^2 = 1$ with the $h_l^2$ parameters giving the proportion of the total random effect variation contributed by each random effect. Our main objective is to conduct inference on these variance component proportions $\boldsymbol{\theta} = [h_1^2,\ldots,h_L^2,h_e^2]$. Secondarily, we may also want to compute the posterior summaries for $\boldsymbol{\beta}$, or for the variance components themselves (i.e. $\sigma_l^2 = \sigma^2 h_l^2$). Therefore, we must calculate the following:

$$p(\boldsymbol{\theta}\,|\,\mathbf{y}) \propto p(\boldsymbol{\theta})p(\mathbf{y}\,|\,\boldsymbol{\theta}) = p(\boldsymbol{\theta})\int_{\boldsymbol{\beta},\boldsymbol{\sigma^2}} p(\mathbf{y}\,|\,\boldsymbol{\beta},\sigma^2,\boldsymbol{\theta})p(\boldsymbol{\beta},\sigma^2)\,d\boldsymbol{\beta}\,d\boldsymbol{\sigma^2} \tag{2}$$

According to Equation 1, the term inside the integral is equal to:

$$\begin{aligned} p(\mathbf{y}\,|\,\boldsymbol{\beta},\sigma^2,\boldsymbol{\theta})p(\boldsymbol{\beta},\sigma^2) = (2\pi\sigma^2)^{-n/2}|\mathbf{V}|^{-1/2} \times \exp\left\{-\frac{1}{2\sigma^2}(\mathbf{y}-\mathbf{X}\boldsymbol{\beta})^\intercal\mathbf{V}^{-1}(\mathbf{y}-\mathbf{X}\boldsymbol{\beta})\right\} \\ \times \frac{b^a}{\Gamma(a)}(2\pi)^{p/2}(\sigma^2)^{p/2+a+1}|\boldsymbol{\Psi}|^{1/2} \times \exp\left\{-\frac{1}{\sigma^2}\left[b + \frac{1}{2}(\boldsymbol{\beta}-\boldsymbol{\mu})^\intercal\boldsymbol{\Psi}^{-1}(\boldsymbol{\beta}-\boldsymbol{\mu})\right]\right\}. \end{aligned} \tag{3}$$

where $\Gamma(\bullet)$ is the gamma function. Using the following linear algebra identity for arbitrary vectors $(\boldsymbol{\gamma},\boldsymbol{\alpha})$ and square matrix $\mathbf{A}$,:

$$\boldsymbol{\gamma}^\intercal\mathbf{A}\boldsymbol{\gamma} - 2\boldsymbol{\alpha}^\intercal\boldsymbol{\gamma} = (\boldsymbol{\gamma}-\boldsymbol{A}^{-1}\boldsymbol{\alpha})^\intercal\mathbf{A}(\boldsymbol{\gamma}-\mathbf{A}^{-1}\boldsymbol{\alpha}) - \boldsymbol{\alpha}^\intercal\mathbf{A}^{-1}\boldsymbol{\alpha},$$

we can rearrange Equation 3 to be:

$$p(\mathbf{y}\,|\,\boldsymbol{\beta},\sigma^2,\boldsymbol{\theta})p(\boldsymbol{\beta},\sigma^2) = C^* \times (\sigma^2)^{-(a+p/2+n/2+1)} \times \exp\left\{-\frac{1}{\sigma^2}\left[b^* + \frac{1}{2}(\boldsymbol{\beta}-\boldsymbol{\mu}^*)^\intercal\boldsymbol{\Psi}^{*-1}(\boldsymbol{\beta}-\boldsymbol{\mu}^*)\right]\right\}, \tag{4}$$

where we define the terms:

$$\begin{aligned} C^* &= \frac{b^a}{\Gamma(a)}(2\pi)^{-(n+p)/2}|\mathbf{V}|^{-1/2}|\boldsymbol{\Psi}|^{-1/2} \\ \boldsymbol{\Psi}^* &= (\boldsymbol{\Psi}^{-1} + \mathbf{X}^\intercal\mathbf{V}^{-1}\mathbf{X})^{-1} \\ \boldsymbol{\mu}^* &= \boldsymbol{\Psi}^*(\boldsymbol{\Psi}^{-1}\boldsymbol{\mu}_{\boldsymbol{\beta}} + \mathbf{X}^\intercal\mathbf{V}^{-1}\mathbf{y}) \\ b^* &= b + \frac{1}{2}\left[\boldsymbol{\mu}^\intercal\boldsymbol{\Psi}^{-1}\boldsymbol{\mu} + \mathbf{y}^\intercal\mathbf{V}^{-1}\mathbf{y} - \boldsymbol{\mu}^{*\intercal}\boldsymbol{\Psi}^{*-1}\boldsymbol{\mu}^*\right]. \end{aligned}$$

Note that if we assume $\boldsymbol{\mu} = \mathbf{0}$, then $b^* = b + \mathrm{RSS}_{\mathbf{V},\boldsymbol{\Psi}}/2$, where $\mathrm{RSS}_{\mathbf{V},\boldsymbol{\Psi}} = \mathbf{y}^\intercal(\mathbf{V}^{-1} - \mathbf{V}^{-1}\widetilde{\mathbf{P}}\mathbf{V}^{-1})\mathbf{y}$ is the (generalized) residual sum of squares given $\mathbf{V}$ and $\boldsymbol{\Psi}$, with $\widetilde{\mathbf{P}} = \mathbf{X}\boldsymbol{\Psi}^*\mathbf{X}^\intercal\mathbf{V}^{-1}\mathbf{X}\boldsymbol{\Psi}^*\mathbf{X}^\intercal$. Equation 4 is the kernel of the NIG$(\boldsymbol{\mu}^*,\boldsymbol{\Psi}^*,a^*,b^*)$ distribution with $a^* = a + n/2$. Therefore, we can evaluate the integral as in Equation 2 as:

$$\int_{\boldsymbol{\beta},\boldsymbol{\sigma}^2} p(\mathbf{y}\,|\,\boldsymbol{\beta},\sigma^2,\boldsymbol{\theta})p(\boldsymbol{\beta},\sigma^2)\,d\boldsymbol{\beta}\,d\boldsymbol{\sigma}^2 = \frac{b^a}{(2\pi)^{(n+p)/2}|\mathbf{V}|^{1/2}|\boldsymbol{\Psi}|^{1/2}\Gamma(a)} \times \frac{(2\pi)^{p/2}|\boldsymbol{\Psi}^*|^{1/2}\Gamma(a^*)}{(b^*)^{a^*}}. \tag{5}$$

In order to evaluate the posterior $p(\boldsymbol{\theta}\,|\,\mathbf{y})$, we must calculate the quantity in Equation 5 for each value in $\boldsymbol{\theta}$ with $p(\boldsymbol{\theta}) > 0$. Using Grid-LMM, we assume this only occurs at a finite (and generally small) set of values on the $L$-dimensional simplex. At each value, we must calculate the determinant of the $n \times n$ dimensional matrix $\mathbf{V}$. However, because of the grid, we efficiently traverse the posterior, reducing the number of evaluations of $\mathbf{V}^{-1}$. All terms on the right hand side of Equation 5 that do not depend on $\boldsymbol{\theta}$ (or $\mathbf{V}$) cancel out in the posterior and Equation 2 can be written in a simplified form as:

$$p(\boldsymbol{\theta} \mid \mathbf{y}) = \frac{|\mathbf{V}_{\boldsymbol{\theta}}|^{-1/2} |\boldsymbol{\Psi}^*_{\boldsymbol{\theta}}|^{1/2} (b^*_{\boldsymbol{\theta}})^{-a^*} p(\boldsymbol{\theta})}{\sum |\boldsymbol{\Psi}_{\boldsymbol{\theta}}|^{-1/2} |\mathbf{V}^*_{\boldsymbol{\theta}}|^{1/2} (b^*_{\boldsymbol{\theta}})^{-a^*} p(\boldsymbol{\theta})} \tag{6}$$

where the subscripts are added to denote each term that is a function of $\boldsymbol{\theta}$. Given this analytical form of $p(\boldsymbol{\theta} \mid \mathbf{y})$, we can now calculate Bayes factors (BF) comparing mixed effect models with different terms in $\mathbf{X}$ [e.g. 1]. In this case, the term $|\boldsymbol{\Psi}|^{1/2}$ will differ among models (i.e. with or without the marker of interest), and therefore will not cancel in the BF ratio. However, as previously shown, if $\boldsymbol{\Psi}$ is diagonal, we can still take the limit of the BF as the hyperparameters $a, b \to 0$ and $\sigma^2_{\boldsymbol{\beta}} \to \infty$ for the covariates (but not for the testing marker which is instead a scale parameter that is specified by the user). This choice of prior distribution class is informative only for the testing SNP, and is otherwise scale-free [1].
